# Deep mutational scanning reveals a shared glycine packing constraint underlying amyloid fibril formation

**DOI:** 10.64898/2026.11.25.769127

**Authors:** Sho Hashimoto, Yoshihisa Watanabe, Teppei Niide, Takashi Nonaka, Kazutaka Katoh, Taro Okunomiya, Nao Harada, Keiko Imamura, Takayuki Kondo, Kayoko Tsukita, Riku Saito, Yuhei Kirita, Mariko Okada, Katsutoshi Taguchi, Masato Hasegawa, Haruhisa Inoue, Satoaki Matoba, Atsushi Hoshino

## Abstract

Amyloid fibrils formed from unrelated proteins adopt diverse folds, yet all consist of identical subunits stacked in register along the fibril axis. Whether this universal architecture imposes common constraints on amino-acid identity has remained unknown. Here, we identify a sequence-independent packing constraint imposed by fibril architecture and use it to predict substitutions that block assembly. Using aggregation-coupled deep mutational scanning, we measured the effects of every amino-acid substitution in α-synuclein on aggregation and abundance. Suppressors clustered at two fibril-core glycines, Gly68 and Gly73, where all substitutions reduced aggregation; G68E and G73C were the strongest human-accessible variants. Structural modeling showed that these residues occupy axial positions where the stacked cross-β lattice cannot accommodate a Cβ atom, with the blocking atom lying in the adjacent layer. G68E restores the ancestral residue present in many mammals. It suppressed aggregation across pathogenic backgrounds, preserved cellular abundance and neuronal localization, and reduced seeded pathology in mouse brain. The constraint is widespread: among 687 fibril folds, 51 of 53 amyloid systems contain at least one buried glycine that cannot accommodate a Cβ atom, usually because of backbone occlusion from the adjacent layer. Applied to amyloid-β, this criterion identified Gly29, and G29E suppressed aggregation in cells. Thus, the architecture that enables amyloid assembly also defines intrinsic vulnerabilities, providing a general principle for identifying candidate protective variants and therapeutic targets upstream of pathology.

## Introduction

Amyloid fibrils form from proteins that share no sequence, structure, or function, yet they assemble into a conserved cross-β architecture in which identical subunits stack in register along the fibril axis (*1*). Cryo-electron microscopy has revealed extensive diversity in these folds, across proteins and across polymorphs of the same protein, while also showing that each fold is reproduced layer by layer along the filament (*2–5*). Despite this structural clarity, the sequence features that a stacked lattice can or cannot accommodate remain undefined. Existing catalogues of amyloidogenic segments (*6*) and of disease-associated mutations (*7–13*) describe protein-specific behavior, but they do not provide a general structural rule that predicts which substitutions a fibril fold inherently excludes. Such a rule would also help identify substitutions of therapeutic interest: variants that block assembly while preserving normal protein function.

α-Synuclein exemplifies this challenge. It is intrinsically disordered, and its assembly into amyloid fibrils is a defining hallmark of Parkinson’s disease and related synucleinopathies (*14, 15*), while the monomer has physiological roles, binding acidic phospholipids and associating with synaptic vesicles (*16–18*). Familial mutations demonstrate that single-residue changes can strongly influence aggregation(*7–12*), yet they represent only a small portion of the possible mutational space. The monomer exists as an ensemble of rapidly interconverting conformations (*14, 19*), so the impact of a substitution cannot be inferred from classical structure-function reasoning. Evolution further complicates the picture: many non-human species, including long-lived mammals, retain the aggregation-prone ancestral Thr53 (*20, 21*) and their α-synucleins fibrillate poorly (*22*), suggesting that other sequence features offset the ancestral residue, features that cannot be identified by surveying human mutations alone.

Comprehensive measurement provides a way to uncover such principles. Using an aggregation-coupled deep mutational scanning (DMS) platform (*23*), we quantified the effects of all 2,641 missense substitutions of α-synuclein on aggregation and abundance, extending earlier mutational surveys of the protein (*19, 24*), and asked what the fibril architecture imposes at the positions the data highlighted. Suppression concentrated at glycines buried in the fibril core, which placed the explanation in the geometry of the stacked lattice rather than in the chemistry of the substituted side chain. We then asked whether the human α-synuclein variant that satisfies this criterion is protective in neurons and in mouse brain. A requirement defined by geometry can be evaluated in any fibril structure, so we finally applied it across the deposited amyloid folds, used it to nominate a substitution at Gly29 in the amyloid-β core (*5, 25*), and tested that substitution in cells.

Because this constraint arises from the assembled state rather than the monomer, substitutions at such positions are expected to inhibit fibril formation while leaving normal protein behavior intact. This is what we observe for G68E, which restores the ancestral residue at position 68 retained by many mammals. It suppressed aggregation on all familial-mutant backgrounds tested and reduced seeded pathology in mouse brain. Localization and presynaptic association were unchanged in knock-in iPSC-derived neurons, as expected for a residue on the cytoplasmic face of the protein rather than at the membrane-binding interface (*18*). These findings reveal a structural requirement shared across amyloid fibrils and show that a naturally occurring human variant already occupies one of the positions at which assembly is most vulnerable to substitution.

## Results

### A subset of human SNCA variants strongly suppresses α□synuclein aggregation

We first constructed a focused human variant mini□library (88 variants) as an initial prioritization of human accessible substitutions, enabling us to identify candidate protective alleles under clinically relevant constraints. The human variant mini library was constructed in the design of C-terminal GFP fusion of α□synuclein (α□synuclein-GFP) protein (Fig. 1A). HEK293T cells transduced with lentivirus to express α□synuclein-GFP form aggregates upon treatment with preformed fibrils (PFF). To selectively detect fibril-formed α□synuclein-GFP, the cells were treated with 0.1% saponin. This mild detergent treatment diffuses non-aggregated α□synuclein-GFP out of the cells and the resulting GFP signal reflects the amount of aggregates present(*26*). This assay allows for the assessment of the aggregation propensity of each cell using FACS (Fig. 1B). Using this saponin-based fibril reporter system, aggregation screening with human SNCA variants mini library was performed. The library-expressing cells were sorted according to the GFP signal and then expanded. Aggregation was induced by treating PFF and incubating for 48 hours. After saponin treatment, cells with and without aggregates were sorted based on the strength of the GFP signal. The read count of each SNCA mutation was analyzed by deep sequencing of extracted genomic DNA (Fig. 1C). To quantify mutation specific changes in aggregation propensity, we defined an aggregation score as the log enrichment of mutant reads in the high aggregation fraction relative to the low aggregation fraction, normalized by the corresponding enrichment observed for the wild-type (WT). This aggregation score was correlated well between replicate experiments: *R^2^* (coefficient of determination) = 0.9504 (Fig. S1A). Among the 88 mutations examined, 48 exhibited a diminished aggregation propensity, while 40 exhibited an augmented property. In the known pathogenic SNCA mutations, H50Q, G51D, A53T, and A53V increased aggregation, while A30P, E46K, and A53E modestly reduced it (Fig. 1D). Because α□synuclein aggregation is central to the pathogenesis of synucleinopathies, variants that strongly suppress aggregation are of particular interest. The top five inhibitory mutations, V52A, G68E, G73C, A90P and A91D, were evaluated individually in both human and mouse SNCA. Mouse WT α□synuclein displayed a high propensity for aggregation due to its possession of the aggregation-prone A53T variant(*19*). All five inhibitory mutations exhibited a strong inhibitory effect on aggregation in both human and mouse SNCA, with G68E demonstrating the most pronounced effect, completely preventing aggregations (Fig. 1E and Fig. S1B and C).

**Fig. 1.**
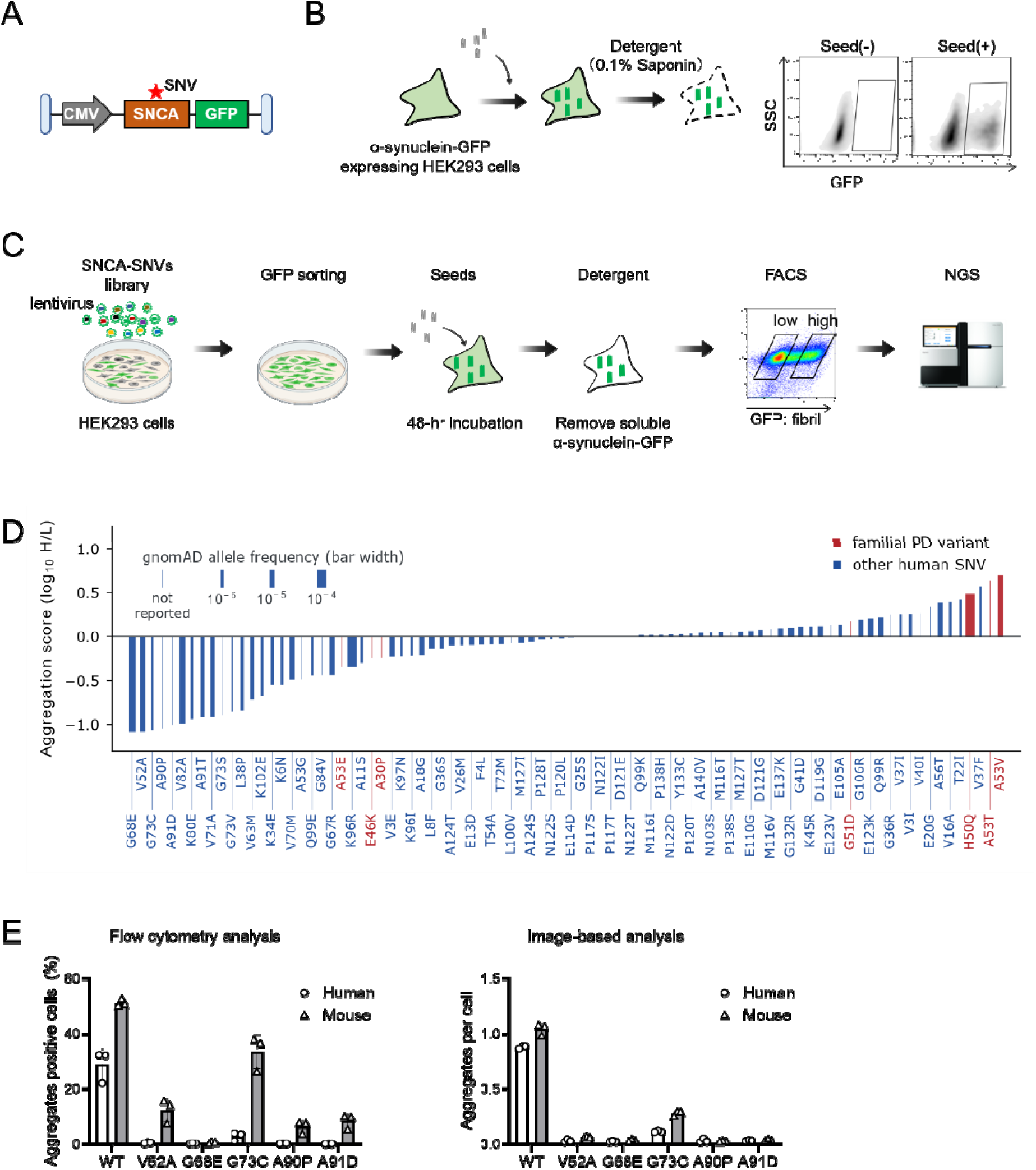
Human SNCA SNVs library screening identifies G68E as the strongest aggregation suppressor. **(A)** Schematic of the α-synuclein–GFP fusion construct used for aggregation-coupled selection. **(B)** Workflow of the saponin-based FACS assay that selectively retains aggregated α-synuclein–GFP. Aggregation was induced by administration of PFF seeds to HEK293T cells expressing α-synuclein–GFP. Treatment with 0.1% saponin passively removed non-aggregated α-synuclein–GFP from the cells, enabling quantification of aggregates by flow cytometry. **(C)** Schematic overview of the screen for α-synuclein aggregation. **(D)** Aggregation scores of 88 human SNCA missense variants, ranked from the most suppressive (left) to the most aggregation-promoting (right). The aggregation score is log□□(H/L), where H and L are the variant counts recovered from the aggregate-high and aggregate-low gates averaged over two independent sorts, expressed relative to WT α-synuclein; negative values therefore denote reduced and positive values increased aggregation. Bar width encodes the allele frequency of each variant in gnomAD v4 (key, upper left); the 18 variants not reported in gnomAD are drawn as hairlines. Reported frequencies span 6.8 × 10□□to 2.6 × 10□□(highest, K96R). Familial Parkinson’s disease variants (A30P, E46K, H50Q, G51D, A53T, A53V, A53E) are shown in red. **(E)** Validation of the five most aggregation-suppressive variants (V52A, G68E, G73C, A90P and A91D) in both human and mouse α-synuclein. Left, flow-cytometric quantification; right, image-based fibril quantification. Data are mean ± SD (n = 3).

### G68E suppresses aggregation across familial mutation backgrounds

We examined the anti-aggregation properties of the G68E variant against pathogenic variants. The G68E variant was introduced into the SNCA gene containing the pathogenic variants in the setting of α□synuclein-GFP fusion protein. Mutated α□synuclein-GFP was expressed in HEK293T cells, and aggregation was facilitated by the administration of PFF. Aggregation was assessed using flow cytometry, which was employed for screening purposes. Additionally, aggregation was quantitatively evaluated using automated imaging and analysis. Consistent with the findings of human SNCA variants screening, the A53T and A53V variants exhibited a pronounced enhancement in aggregation, while the G51D exhibited a modest aggregation propensity. Conversely, the A30P, E46K and A53E variants demonstrated a reduction in fibril generation. In the background of these pathogenic variants, the G68E variant still completely prevented aggregation (Fig. 2A).

**Fig. 2.**
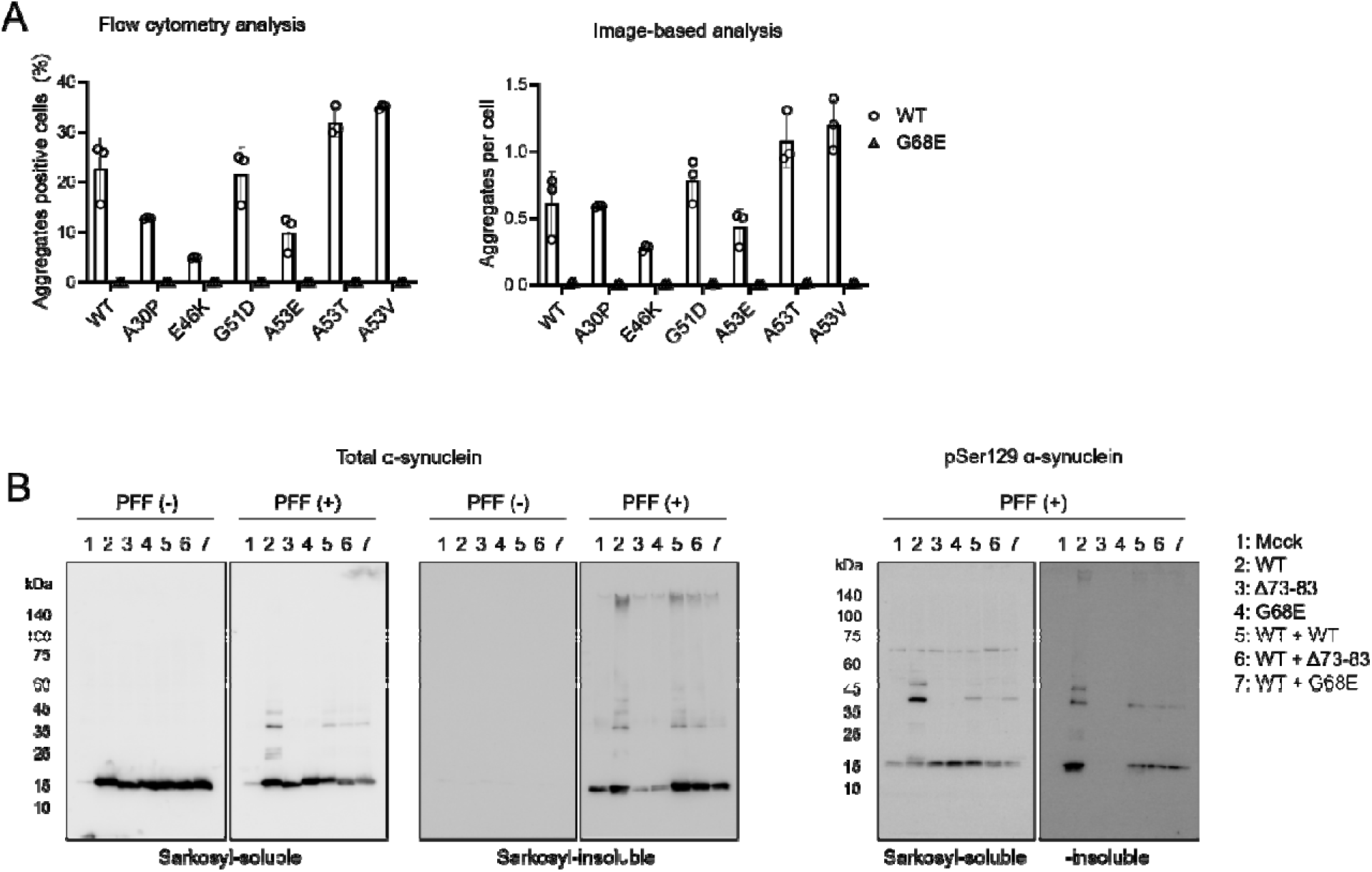
G68E variant has the capacity to impede fibril and oligomer formation. **(A)** G68E-mediated inhibition of aggregation was observed in the background of pathogenic variants. The left panel is the flow cytometry analysis, and the right panel is the image-based fibril quantification. Data are presented as mean ± SD (n = 3). **(B)** The anti-aggregation effect of G68E α□synuclein against WT α□synuclein was evaluated in comparison with the core domain-deleted Δ73-83 α□synuclein. HEK293 cells expressing the constructs listed at the right were left untreated (PFF −) or given α-synuclein pre-formed fibrils (PFF +), lysed in buffer containing 1% Sarkosyl, and the supernatant (Sarkosyl-soluble) and pellet (Sarkosyl-insoluble) fractions analyzed by western blotting. Lane 5 (WT + WT) is the expression-matched control for lanes 6 and 7. The pSer129 blots were obtained for the PFF (+) condition only. Representative of n = 4 independent experiments.

β-Synuclein, which lacks the hydrophobic non-amyloid-β component (NAC) region, has been reported to inhibit the nucleation and elongation of α-synuclein fibrils by competing for membrane and fibril surfaces (*27, 28*). We next examined whether G68E acts in the same way. WT or G68E α-synuclein, alone or co-expressed with WT, was seeded with PFF in HEK293 cells, and the sarkosyl-soluble and sarkosyl-insoluble fractions were examined by western blotting, with the Δ73–83 mutant, a β-synuclein-like deletion that removes the central NAC segment, as a comparator (Fig. 2B). Expressed alone, neither G68E nor Δ73–83 gave pS129-positive insoluble α-synuclein, as in mock-transfected cells. Whenever WT α-synuclein was present, however, that signal was readily detected, and neither construct reduced it relative to the WT + WT control, although soluble α-synuclein was comparable in the WT + WT and WT + G68E conditions. Because the recombinant seeds carry no pS129, the signal reports aggregates formed from cellular protein. G68E therefore suppresses only its own recruitment, and the trans-inhibition reported for β-synuclein was not reproduced by its NAC-deleted mimic.

### Deep mutational scanning reveals the global aggregation landscape of α□synuclein

To comprehensively understand the effects of mutations on the aggregation propensity of α□synuclein, DMS was conducted with the library containing all single amino acid substitutions of human α□synuclein. The DMS library was constructed as an α□synuclein-GFP fusion followed by an internal ribosomal entry site (IRES) and miRFP670 as a marker of construct expression (Fig. S2A). The library was transduced into HEK293T cells, and cells expressing the constructs were isolated by sorting for the miRFP670 signal. To analyze the aggregation properties, the library cells were treated with PFF and sorted according to the GFP fibril signal after saponin treatment (Fig. S2B). In addition, the impact on protein abundance was assessed. Both the high and low GFP signal populations, relative to the miRFP670 signal, were sorted (Fig. S2B). The abundance score was calculated as the log enrichment of mutant reads in the high-GFP fraction relative to the low□GFP fraction, normalized by the corresponding enrichment observed for the WT. Aggregation scores were highly reproducible between replicates (*R*² = 0.8738), whereas abundance scores were moderately reproducible (0.5621) (Fig. S2C). The library comprised 2,641 missense substitutions and 139 nonsense variants across residues 2 to 140 (Fig. 3A). Because the GFP reporter lies C-terminal to α□synuclein, a premature stop abolishes the fusion protein itself: the nonsense variants collapsed in both assays (aggregation −0.83 ± 0.17, abundance −1.04 ± 0.37), confirming the dynamic range of the screen (Fig. 3B).

**Fig. 3.**
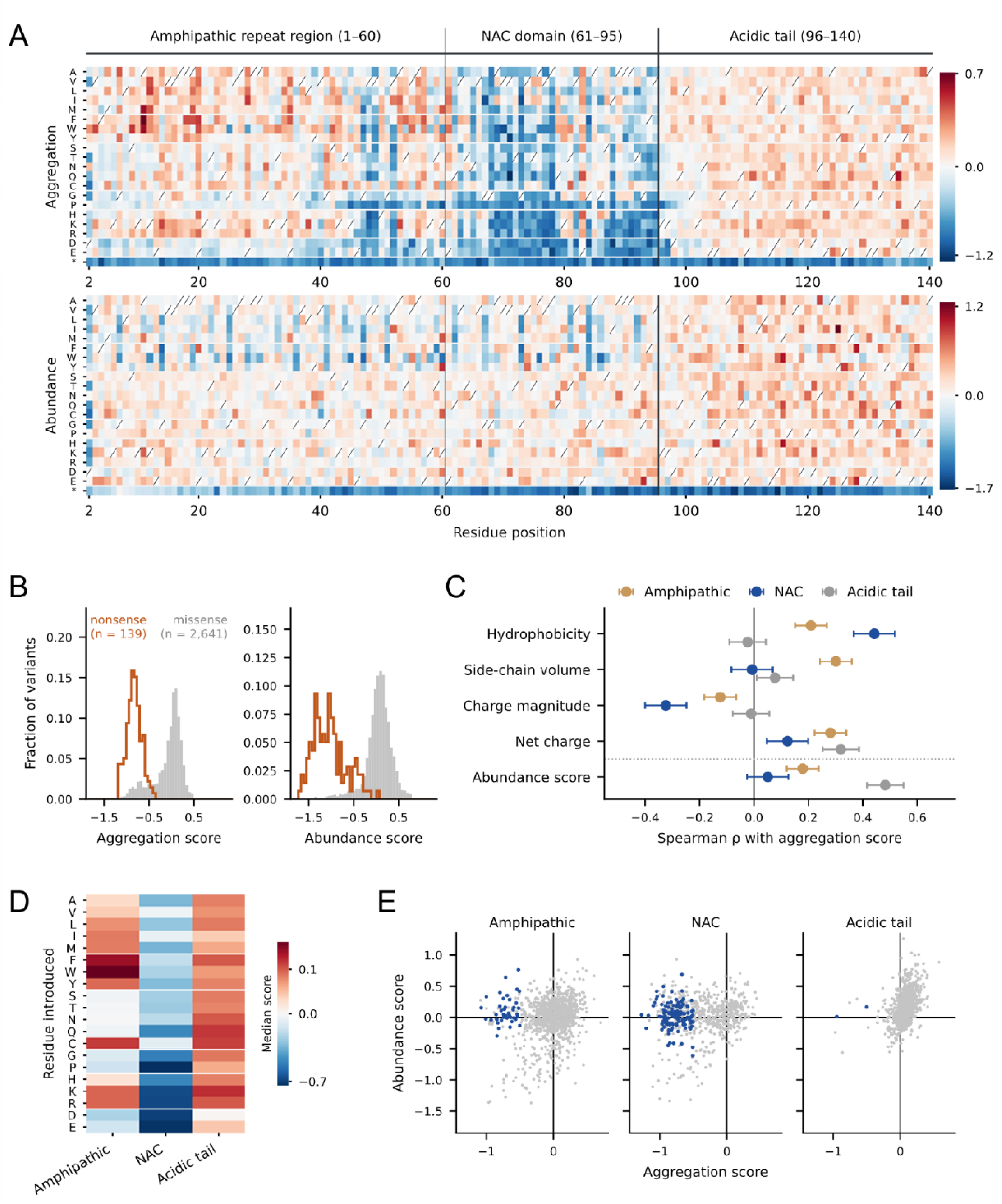
Deep mutational scanning of α-synuclein aggregation and abundance. **(A)** Variant-effect maps for aggregation (top) and protein abundance (bottom). Each column is one of the 139 positions assayed (residues 2 to 140) and each row one of the 20 amino acids or the stop codon; rows are grouped as aliphatic, aromatic, polar, glycine and proline, basic and acidic. Color gives the score according to the scale bar on the right and the WT residue at each position is crossed by a diagonal line. **(B)** Distribution of the aggregation score (left) and the abundance score (right) for all missense substitutions (gray) and for the 139 nonsense variants (orange), each histogram normalized to the number of variants in its own group. Because the GFP reporter lies C-terminal to α-synuclein, a premature stop codon removes the fusion protein itself; the nonsense variants (aggregation −0.83 ± 0.17, abundance −1.04 ± 0.37) therefore mark the floor of each assay. **(C)** Spearman correlation between the aggregation score and four properties of the substitution: the change in side-chain hydrophobicity on the Kyte–Doolittle scale, the change in side-chain volume, the magnitude of the charge introduced and the signed change in net charge. Below the dotted rule, the correlation with the abundance score measured in the same library. Correlations are computed separately within each region (amphipathic repeat region, residues 2 to 60, n = 1,121; NAC domain, residues 61 to 95, n = 665; acidic tail, residues 96 to 140, n = 855); error bars, the 95% confidence half-width for that number of substitutions. **(D)** Median aggregation score of substitutions grouped by the residue introduced (rows) and the region substituted (columns, as in c), on its own diverging scale. **(E)** Aggregation against abundance score for all 2,641 substitutions, separated by region as in c. Blue, the 182 substitutions that suppress aggregation (q < 0.05 and score below −0.5) while their abundance score remains statistically indistinguishable from WT given the replicate error (|score| < 1.96 × s.e.m.).

The maps show a clear regional organization in both assays (Fig. 3A). In the amphipathic repeat region the aggregation score scaled with the residue introduced, correlating with the change in side-chain volume (Spearman ρ = +0.30), in hydrophobicity (+0.21) and in net charge (+0.28, and +0.31 with the abundance score held constant; Fig. 3C): hydrophobic or positively charged substitutions increased aggregation, whereas negatively charged or helix□disrupting proline substitutions reduced it (Fig. 3D). Physiologically, α□synuclein specifically binds to anionic phospholipids, such as phosphatidylserine (PS) through the electrostatic interaction between the positively charged lysine residues in the N-terminal region and the negative headgroups of the lipids (*29, 30*), and membrane binding converts the region into an amphipathic α helix that associates with curved, acidic synaptic vesicles (*31*). The NAC region, dominated by alanine and valine, was far more sensitive, and what mattered there was the chemistry of the side chain rather than its size: the score correlated with hydrophobicity (ρ = +0.44) and with the magnitude of the charge introduced (−0.32), but not with side-chain volume (−0.01) or with whether that charge was positive or negative (+0.12; Fig. 3C). Charged residues were accordingly the strongest side-chain suppressors, glutamate (median −0.73), aspartate (−0.70), arginine (−0.68) and lysine (−0.67), as expected where an in-register stack repeats the same charge in every layer, adding electrostatic repulsion along the fibril axis and, at buried positions, a desolvation cost (Fig. 3D). Consistent with the repulsion term, substitutions at Glu61 and Glu83, the two negatively charged residues of the region, promoted aggregation (medians +0.19 and +0.25, against −0.39 for the region as a whole; Fig. 3A). Proline suppressed more strongly still (−0.75), but through the backbone rather than the side chain: its rigid cyclic structure and absence of an amide hydrogen are incompatible with the β-strand. The acidic tail responded to neither hydrophobicity nor volume (ρ = −0.02 and +0.08); the only property that mattered was the loss of negative charge, which increased aggregation (+0.32, and +0.33 with the abundance score held constant; Fig. 3C), as expected if the tail restrains NAC-driven assembly by electrostatic shielding.

Considering the disease-protective variants, substitutions that reduce aggregation while preserving abundance could be candidates if they maintain normal physiological function. Of the 2,641 substitutions, 182 suppressed aggregation (q < 0.05 and score below −0.5) while leaving the abundance score indistinguishable from the WT, 21 of them at glycine positions. Substitutions of this class were concentrated in the NAC domain: of the 182, 138 lay in residues 61 to 95, 42 in the amphipathic region and 2 in the acidic tail (Fig. 3E). Among the 134 substitutions with the strongest suppression (below −0.8) only 22% showed reduced abundance, so aggregate suppression was generally not a consequence of losing the protein.

### Two core glycines occupy positions at which the fibril lattice cannot accommodate a Cβ atom

The 182 substitutions that suppressed aggregation without reducing abundance came from positions of two kinds. At four buried aliphatic residues, Val49, Val71, Val77 and Ala78, all 19 substitutions suppressed aggregation, as expected where substitution removes a side chain from the hydrophobic core. The only other positions with that uniform intolerance were the glycines Gly73 and Gly68, ranked first and fourth of the 139 positions assayed (mean of the 19 substitutions, −0.81 and −0.70); there suppression was independent of charge, polarity and volume, and did not reflect reduced abundance. The effect extends to the core glycines as a group: their positional means average −0.40, against −0.07 for the 121 non-glycine positions, whereas the nine glycines that lie outside the ordered core average +0.07 (Fig. 4A and B). A glycine has no side chain to remove: what its 19 substitutions share is the addition of a Cβ atom where glycine carries only a hydrogen. To test whether this atom is sterically incompatible with the fibril, we modeled a Cβ at each of the nine core glycines and measured the free radius, the largest sphere that fits without overlapping a neighboring atom. In the three multiple system atrophy filaments (PDB 6XYO, 6XYP, 6XYQ) (*2*), no position reaches the carbon radius, but the atom responsible differs. At six of the nine it belongs to the same subunit, where a side chain can move aside. At the other three, Gly68, Gly73 and Gly84, it belongs to the adjacent subunit along the fibril axis, one layer up or down. Measured against the adjacent subunit alone, these three are the tightest of the nine glycines in all three filaments (0.44 to 1.24 Å; Fig. 4C and Fig. S3A). Only Gly68 and Gly73, however, remain below the carbon radius in all six structures (Fig. 4C). That axial clash can be relieved only by breaking the hydrogen bonds that hold the layers together. In 6XYO the stack would have to open from its observed 4.73 Å rise to 6.49 Å before a carbon fits at Gly68, and the inter-subunit backbone hydrogen bonds do not survive that expansion: the first breaks at 5.11 Å and all 137 are gone by 5.74 Å (Fig. 4D and E). A lattice with room for a Cβ at position 68 is no longer held together. The same measurement at Gly73 gives 0.56 Å in 6XYO, and the position is below the carbon radius in all six structures (0.56 to 1.34 Å; Fig. S3B). There the hydrogen-bond argument is weaker because less expansion is needed: reaching the carbon radius requires a rise of 5.15 to 6.19 Å, which breaks every inter-subunit backbone hydrogen bond in 6XYO, 6XYQ and 8A9L but only 51% of them in 6XYP, 26% in 6H6B and 5% in 6SSX (Fig. S3C).

**Fig. 4.**
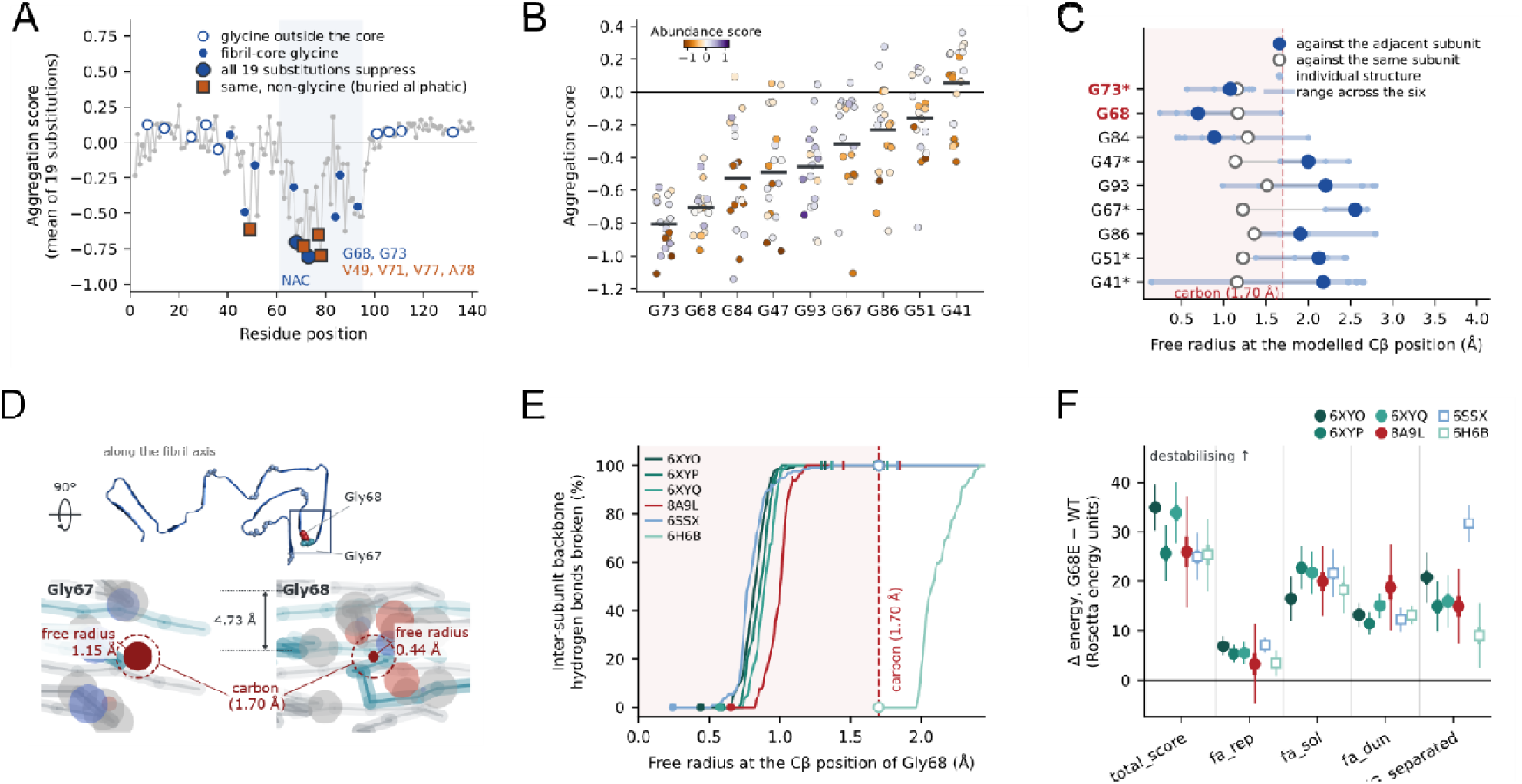
Steric blockade of the Cβ position, not backbone conformation, marks the suppressive glycines. **(A)** Mean aggregation score of the 19 substitutions at each of the 139 positions assayed (gray), with the nine glycines of the ordered core (filled blue; mean −0.40), the nine outside it (open blue; +0.07) and the remaining positions (−0.07); band, NAC region (residues 61 to 95; −0.38 within against +0.02 outside). The core glycines differ from the remaining 121 positions (two-sided Mann-Whitney P = 8 × 10□□). Ringed symbols, the six positions at which all 19 substitutions suppressed aggregation (Gly68, Gly73, Val49, Val71, Val77, Ala78). **(B)** All 19 substitutions at each glycine, ordered by positional mean (bars); each point one substitution, colored by abundance score. **(C)** Free radius at the modeled Cβ position of each core glycine, against atoms of the same subunit (open) and of the adjacent subunits along the fibril axis (filled, mean of the six structures; small points, individual structures; bar, their range), in three multiple system atrophy filaments (PDB 6XYO, 6XYP, 6XYQ), one Lewy-fold filament (8A9L) and two polymorphs assembled in vitro (6SSX, 6H6B). On the mean, Gly68, Gly73 and Gly84 (0.70 to 1.08 Å) fall left of the carbon radius (dashed, 1.70 Å) and the six tolerant positions (1.91 to 2.56 Å) right of it; only Gly68 and Gly73 (red) lie left of it in every structure. Asterisks, left-handed backbone (φ > 0) in the brain filaments. **(D)** Top, the 6XYO subunit viewed along the fibril axis, with the nine core glycines as spheres (Gly68 red, Gly67 teal); the box marks the region drawn below and the glyph the 90° rotation between the two views. Bottom, lattice cross-section at the modeled Cβ position of Gly67 and Gly68, in the plane containing the fibril axis (vertical) and the Cα-Cβ direction: discs, van der Waals spheres cut by that plane (carbon gray, nitrogen blue, oxygen red); bands, Cα traces of the central subunit (teal) and its two neighbors (gray), fading with distance from the plane; dark red disc, the sphere that fits (1.15 Å at Gly67, 0.44 Å at Gly68); dashed circle, carbon; arrow, the axial rise per subunit (4.73 Å). At Gly67 the space is bounded by the same subunit, at Gly68 by the subunit above (Asn65 ND2, Gly68 O, Asn65 CG). **(E)** Cost of making room at Gly68. With the stack expanded rigidly along the fibril axis, the percentage of inter-subunit backbone hydrogen bonds exceeding 3.5 Å (n = 57 to 137 per structure) against the free radius reached. Filled circles, observed rise; ticks, rises of 5.5, 6.0 and 6.5 Å; open circles, where axial expansion relieves the adjacent-subunit clash (rise 4.93 to 6.57 Å). **(F)** Rosetta energy difference (G68E − WT) per term in all six structures. Symbols, difference between the means of 50 independently relaxed models per condition; bars, the 95% confidence interval of that difference (thick) inside its standard deviation (thin). All terms differ from zero in every structure (P < 0.005, Welch’s t test).

Glycine also samples backbone conformations that are rare for other residues, so the intolerance could in principle be conformational rather than steric: the position might need a conformation only glycine can take (*32*). The reference distributions rule that out. At Gly68, every one of the 18 non-proline residue types is observed within 30° of the conformation the position adopts, and this holds in all six structures. Across the nine core glycines, 48 of the 54 position-structure conformations fall inside the region where all 18 types are observed, proline being the only type the analysis excludes. A conformational requirement would also have to hold across folds, yet at most of the nine positions the conformation differs between them (Fig. S3, D and E). The distinguishing feature of Gly68 and Gly73 is therefore steric: the space available to a Cβ atom, not the φ/ψ angles of the position, which every residue type but proline can adopt. The steric criterion is nonetheless fold-specific (Fig. S3A). Gly68 is the tightest of the nine in the in vitro polymorph 6SSX (0.24 Å) (*3*) and remains blocked in the Lewy fold 8A9L (0.65 Å) (*33*), where only Gly41 is tighter, whereas in 6H6B (*4*), Gly68 is blocked by the narrowest margin of the six (1.68 Å) and an expansion of 0.02 Å admits a carbon. The cellular assay is seeded with recombinant preformed fibrils, and which of these folds it templates is not known.

We took G68E forward, the strongest suppressor in the human single-nucleotide-variant screen (Fig. 1D). It destabilized all six structures by +25.0 to +34.9 Rosetta energy units (Fig. 4F). The repulsive term contributes least (fa_rep, +3.3 to +7.1); most of the cost is desolvation (fa_sol, +16.5 to +22.8) and side-chain rotamer strain (fa_dun, +11.4 to +18.8), with a further loss of inter-chain binding energy (dG_separated, +9.1 to +31.7). Relaxation cannot remove the clash, only convert it into strained rotamers that bury the glutamate.

### Glu68 and Gly95 as protective states tolerating the ancestral aggregation-prone Thr53

Although the A53T mutation in α□synuclein causes early□onset Parkinson’s disease in humans, threonine at position 53 is the WT residue in many other species, including New World monkeys. Yet these species, even long□lived ones, show no α□synuclein pathology(*20, 34*), implying the presence of additional protective factors that prevent disease despite aggregation-prone Thr53. Evolutionary conservation analysis of α□synuclein in vertebrates revealed that highly conserved residues cluster within the N□terminal amphipathic region containing the KTKEGV repeats, whereas the central NAC region shows strong preservation of hydrophobic residues associated with fibrillogenic propensity (Fig. 5A and S4). In contrast, the acidic C terminal tail displays lower conservation, reflecting its role as a flexible, regulatory domain that remains structurally disordered (Fig. 5A), suppresses NAC□driven aggregation through electrostatic shielding, and serves as a hub for protein interactions and post□translational modifications(*35, 36*). This profile highlights evolutionary constraints on membrane□binding and aggregation□modulating elements of α□synuclein. Integrating evolutionary conservation with our DMS results identified G68E and V95G as candidates to suppress the aggregation propensity, even in the presence of the aggregation-prone A53T. Mapping residues 53, 68, and 95 onto a vertebrate phylogeny highlights how different segments of α-synuclein have been shaped by distinct evolutionary pressures (Fig. 5B). Ancestral reconstruction indicates that Thr53, Gly68, and Val95 represent the ancestral states of vertebrate α□synuclein. While Ala53 marks the divergence of Old World and New World primates as previously reported(*21*), Glu68 arose once, on the stem of placental mammals, and is the ancestral placental state: across 69 placental mammals maximum-parsimony reconstruction requires only a single Glu-to-Gly loss on the stem of Euarchontoglires and a single reversal in *Oryctolagus cuniculus* (Fig. S5A); in the 169-species vertebrate tree the only other independent Gly-to-Glu gains are in four teleost lineages. Residue 95 shows more relaxed constraints, with the tuatara independently acquiring Gly95 despite its absence in most mammals. Because Glu68 is ancestral in placental mammals rather than repeatedly acquired, its distribution is not in itself evidence of selection in long-lived lineages, and Glu68 species are not longer-lived than Gly68 species after correction for phylogeny and body mass (Fig. S5, B and C). What the distribution does establish is tolerance: whole placental lineages carry Glu68 in a protein that must still function. We then tested the residues directly, in seven long-lived species spanning three of the four combinations of residues 68 and 95. The bowhead whale (*Balaena mysticetus*) and the African elephant (*Loxodonta africana*) have both G68E and V95G, while the tuatara (*Sphenodon punctatus*) and the red-fronted lemur (*Eulemur rufifrons*) have only V95G. In contrast, the Pinta Island tortoise (*Chelonoidis abingdonii*), the naked mole-rat (*Heterocephalus glaber*), and the Komodo dragon (*Varanus komodoensis*) have neither (Fig. S4). The aggregation properties of α-synuclein were closely linked to the presence of the Glu68 and Gly95. Specifically, aggregation was inhibited in the bowhead whale and African elephant, while the tuatara and red-fronted lemur exhibited modest aggregation. Other species showed aggregation levels comparable to those of humans and mice (Fig. 5C). Converting Glu68 and Gly95 to the human residues Gly68 and Val95 raised aggregation to the level of human α-synuclein (Fig. 5D). These results indicate that naturally occurring substitutions such as G68E and, potentially, V95G can act as aggregation-suppressive modifiers that counteract the intrinsic aggregation propensity of α□synuclein, enabling certain lineages to tolerate the otherwise pathogenic ancestral Thr53 background.

**Fig. 5.**
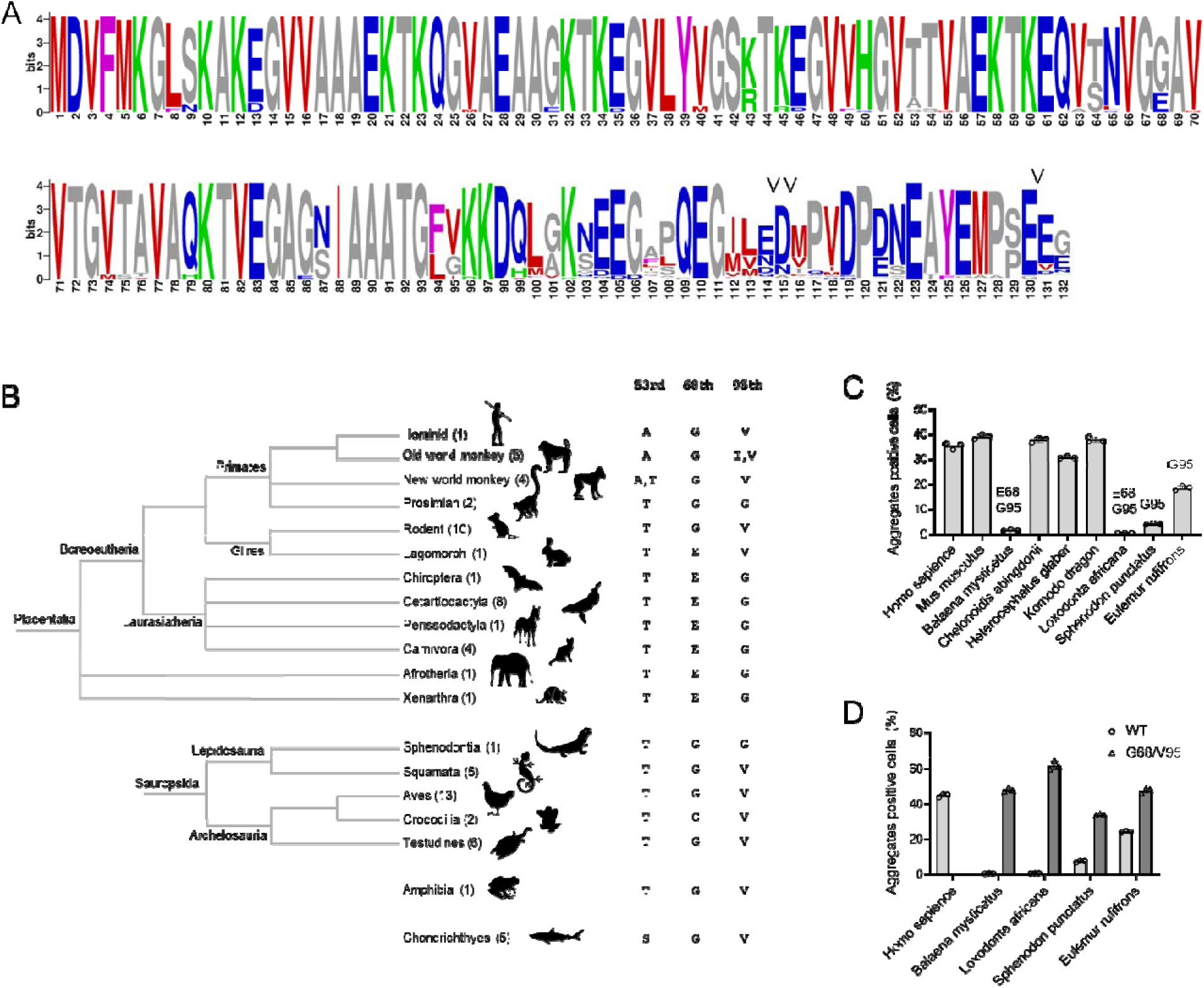
Natural Glu68 and Gly95 states act as protective modifiers against α-synuclein aggregation. **(A)** Sequence logo representation of α□synuclein conservation across vertebrate species. A multiple sequence alignment of representative vertebrate α□synuclein orthologs was used to compute residue wise conservation, shown as information content (bits) on the y□axis. The C□terminal region was excluded from the analysis due to uncertainties in its sequence alignment. The height of each amino acid letter reflects its relative frequency at that position, and colors denote physicochemical classes. V indicates insertion in some species. **(B)** Cladogram mapping of residues 53, 68, and 95 across vertebrates. Numbers in parentheses indicate the number of species represented in each clade. **(C)** The aggregation propensity of long-lived animals, some of which contain G68E and/or V95G, displays limited aggregation. Data are presented as mean ± SD (n = 3). **(D)** The aggregation propensity of conversion to human G68 and/or V95 in the animals containing G68E and/or V95G α□synuclein. Data are presented as mean ± SD (n = 3).

### G68E suppresses seeded α-synuclein pathology without altering normal protein localization

To assess the subcellular localization of the G68E variant, we generated homozygous G68E knock□in iPSCs and performed immunocytochemistry in iPSC-derived neurons. Homozygous G68E knock-in neurons displayed α-synuclein localization comparable to WT protein. The distribution of α-synuclein was observed in the soma, neurites, nucleus, and synaptic sites, where it colocalized distinctly with synaptic markers, such as Synapsin-1 (Fig. 6A). α-Synuclein binds membranes through an amphipathic helix formed by its N-terminal 11-mer repeats, and residue 68 lies at the fourth register position, where six of the seven other repeats carry glutamate or glutamine (Fig. S6A). Placed in an implicit bilayer, residues 61 to 84 sit nearly parallel to the membrane with only the hydrophobic face inserted: Val63 reaches 3.7 Å past the hydrocarbon-core boundary while Gly68 lies 0.3 Å outside it (Fig. S6B to D). The calculation also predicts that membrane binding is unaffected, in line with the localization we observe.

**Fig. 6.**
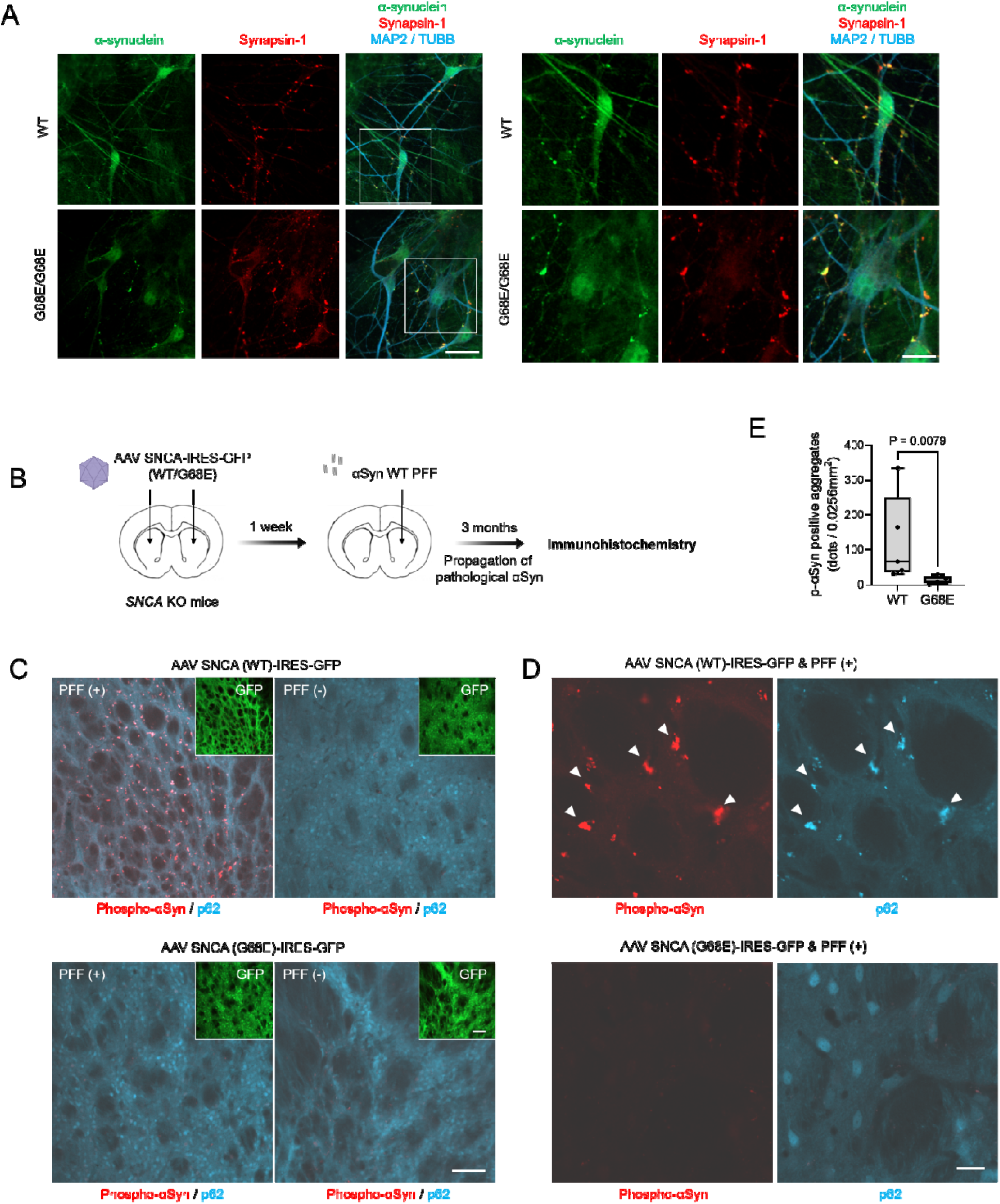
G68E prevents fibril propagation in the mouse brain. **(A)** Immunofluorescence analysis of α□synuclein distribution in WT and homozygous G68E iPSC-derived neurons. Neurons were stained for α synuclein (green), Synapsin□1 (red), and MAP2/TUBB (cyan) to visualize presynaptic sites and neuronal morphology. Representative fields (left) and magnified views of boxed regions (right; panels a–c) demonstrate that α□synuclein exhibits a punctate distribution along neurites and robust colocalization with Synapsin□1–positive presynaptic boutons in both genotypes. Arrowheads indicate representative colocalized puncta. No overt differences in the pattern or extent of synaptic enrichment were observed between WT and G68E mutant neurons. Scale bars, 20 μm (left) and 10 μm (right). **(B)** Workflow of AAV-mediated α synuclein expression in the brain of SNCA knockout mice and evaluation of fibril propagation. AAVs carrying genes of WT or G68E SNCA-IRES-GFP were stereotactically injected into the bilateral striatum, and one week later, PFF was injected into the striatum on one side. **(C)** Immunohistochemistry of the brain expressing AAV SNCA WT or G68E-IRES-GFP, with or without α□synuclein PFF treatment, showing images of phospho□α□synuclein (red), p62 (blue), and GFP (green) for each genotype and condition. Robust phosphor-α□synuclein- and p62-positive aggregates are detected exclusively in WT neurons following PFF exposure. Scale bar, 100 μm. **(D)** Magnified individual images for phospho-α-synuclein and p62 in PFF-treated brains, with arrowheads marking representative puncta where phospho α□synuclein and p62 signals colocalize. Scale bar, 20 μm. **(E)** The α□synuclein aggregates were quantified as phosphorylated α□synuclein dots per unit area. Data are presented as individual data points overlaid on box and whisker plots (n = 5 per group). Statistical analyses were performed using unpaired two-tailed Mann-Whitney test.

We further evaluated the impact of G68E on pathological propagation of α□synuclein aggregates *in vivo*. AAVs carrying either human WT or G68E mutant SNCA, with IRES-GFP as a transduction marker, were injected into the bilateral striatum of 8-week-old SNCA□knockout mice. One week later, PFF seeds were administered into the striatum on one side. Three months after fibril inoculation, brain tissues were collected and subjected to histological analysis (Fig. 6B). The expression of α□synuclein was detected together with GFP fluorescence, which confirmed equivalent AAV expression between WT and G68E, excluding expression-level effects (Fig. S7). Mice expressing WT α□synuclein developed abundant phosphorylated α□synuclein aggregates that colocalized with p62, which is a feature of Lewy body pathology. In contrast, those expressing the G68E variant showed a marked reduction in aggregate formation (Fig. 6C to E). Collectively, these findings suggest that G68E suppresses α□synuclein aggregation and may function as an aggregation-suppressive variant.

### A glycine packing constraint shared by amyloid folds

The positions identified in α-synuclein are set by the geometry of the lattice rather than by the sequence, so the same measurement can be made in any fibril fold. Across 687 non-redundant fibril folds of 53 amyloid systems the constraint is widely shared: 70.1% of the 4,697 buried core glycines are blocked by an atom in an adjacent layer, against 7.5% of the 29,807 buried non-glycines. Fifty-one of the 53 systems contain at least one glycine blocked by an adjacent layer. Of the remaining two, each of which offers a single glycine in its core, one (glucagon) is blocked instead by an atom of the same subunit and the other (NLRP6) is not blocked at all. The proportion of a system’s core glycines that are blocked by an adjacent layer has a median of 71% (interquartile range 60 to 78%), reaching 100% in five systems and 0% only in the two whose core contains a single glycine; if glycines were blocked independently at the pooled rate, 1.7 systems would be expected to contain none (Fig. 7A and Fig. S8A and B). The existence of the constraint is not amyloid-specific. Among 21 natively folded proteins, 91.3% of the 92 buried glycines are blocked as well, and the free radius left at their Cβ position is distributed as it is in the fibril cores (medians 1.03 and 1.03 Å; Fig. S8B). What differs is the source of the block. In a globular fold the blocking atom belongs to the same chain; in a fibril it lies one layer away. Most often it is the cross-β backbone itself: of the 3,294 axially blocked glycines, a backbone atom is the limiting atom in 70.3%, and in 50.6% that atom is the carbonyl oxygen of the glycine’s own counterpart in the next layer (Fig. S8C). Among the 104 glycines measurable both in the native fold and in the fibril of the same protein (13 proteins, 95 folds), 46 positions that accommodate a Cβ while the protein is folded cease to accommodate one in the fibril, against 4 that do the reverse, and the median free radius falls from 1.85 Å to 0.95 Å (Fig. 7B). The shift is neither uniform nor purely axial: the losses concentrate in lysozyme and transthyretin (22 of 46), and 52 of the 104 glycines were buried natively, so part of the change reflects burial on assembly rather than the stack alone.

**Fig. 7.**
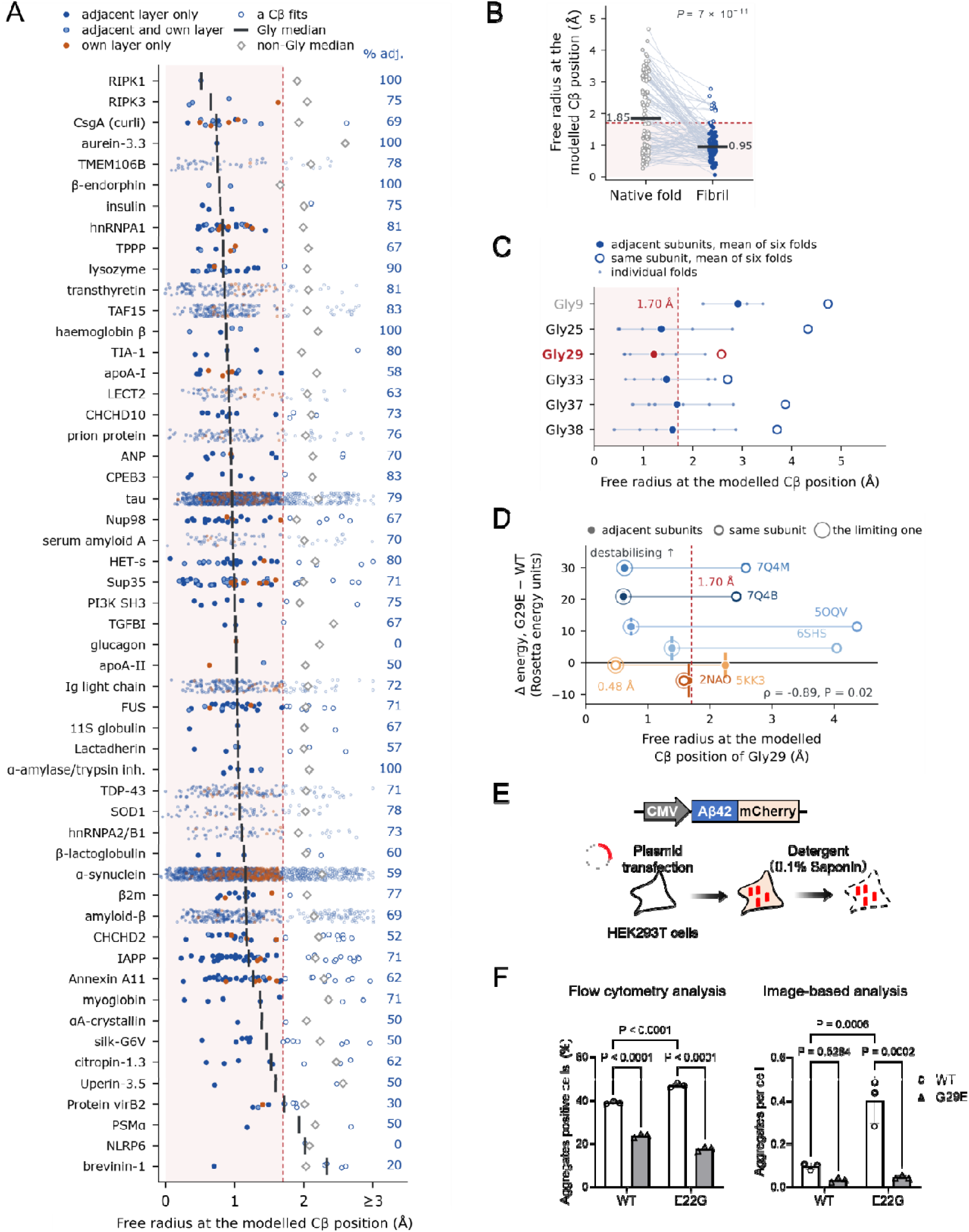
The Cβ packing constraint is a general consequence of cross-β assembly and operates in amyloid-β. **(A)** Free radius at the modeled Cβ position of every buried core glycine, by amyloid system: 4,697 positions in the 53 systems that have one, from 687 non-redundant fibril folds, ordered by median with the most constrained at the top. Buried is a relative solvent accessibility below 0.25; each position of a fold contributes one observation; a system merges entries of the same protein across species, isoforms and point-mutant constructs, so the 53 systems correspond to 70 UniProt entries. Color gives the origin of the blockage, the black bar is the median of the system, and the gray diamond the median of the buried non-glycines of the same cores, at which a Cβ is in fact present. Shading and the dashed line mark the carbon radius; 11 positions lie beyond the axis and are drawn at its right edge. % adj., percentage blocked from an adjacent layer. **(B)** Free radius for the 104 glycines measurable in both the native fold and the fibril of the same protein (13 proteins, 95 non-redundant folds); lines join the same glycine in the two states, thick bars are medians, filled markers the 91 positions blocked in the fibril. The fibril value is the median over the folds of that protein. The median falls from 1.85 to 0.95 Å (Wilcoxon signed-rank P = 7.1 × 10-11). **(C)** Free radius at the modeled Cβ of each ordered-core glycine of amyloid-β, against atoms of the same subunit (open) and of the adjacent subunits along the fibril axis (filled, mean of six folds; small points, individual folds; bar, range). Folds: 7Q4B and 7Q4M (Alzheimer brain), 5OQV (Aβ42 in vitro) and 6SHS (Alzheimer brain, Aβ40) by cryo-electron microscopy, 2NAO and 5KK3 by NMR. Dashed line, carbon radius (1.70 Å). On the adjacent-subunit measure Gly29 (red) is the only position below 1.70 Å in every fold; Gly9 (gray) is ordered in four folds and buried in none. **(D)** For each fold, the free radius left at the modeled Cβ position of Gly29 by the adjacent subunits (filled) and by the same subunit (open), joined by a line, against the Rosetta repacking penalty of G29E (Δ total score, G29E minus WT, mean of 50 relaxed models per state, 95% confidence interval on the filled marker). The ring marks the limiting measure and the printed value the tightest position in the panel; dashed line, carbon radius. In the four cryo-electron microscopy folds the adjacent subunits set the limit while the same subunit leaves 2.4 to 4.4 Å; in the two NMR folds the order reverses and the limiting atom is an Ile41 side chain within the subunit. The penalty scales with the adjacent-subunit free radius (Spearman ρ = −0.89, P = 0.02, on the filled markers). Decomposition into energy terms in Fig. S8D. **(E)** A saponin-based assay to quantify intracellular Aβ aggregates. Overexpression of Aβ42-mCherry in cells led to the formation of intracellular aggregates even in the absence of exogenously added PFF seeds. **(F)** Aggregate content, WT against G29E, on the WT and the pathogenic E22G background, by flow cytometry and by image analysis.

We next tested the criterion in amyloid-β. Its fibril core contains five glycines (Gly25, Gly29, Gly33, Gly37 and Gly38; Gly9 lies outside it). Among six fibril folds, two from Alzheimer brain (*5, 37*), two assembled in vitro(*38*) and two determined by NMR (*39, 40*), Gly29 has the smallest median free radius (0.68 Å, against 1.17 to 1.51 Å at the other four glycines) and is the only one that falls below the carbon radius in every fold (Fig. 7C). In the four cryo-EM structures, including both brain-derived folds, Gly29 is bounded axially by the adjacent subunit, as α synuclein Gly68 is; in the two NMR folds the limiting atom lies within the layer. That distinction sets the size of the energetic penalty, not merely its presence. The repacking penalty for G29E tracks how closely the adjacent layer approaches the Cβ position, and it disappears in the two NMR models, the two folds in which that layer sits furthest away (Fig. 7D). In 5KK3, the Cβ position is more crowded than in any of the four folds where G29E is destabilizing (0.48 Å against 0.60 to 1.39 Å), yet the substitution costs nothing there, because the limiting atom is a same-layer side chain that can move aside (Fig. 7D and Fig. S8D). In cells expressing Aβ42-mCherry, G29E reduced aggregation on the WT background and even on the pathogenic E22G background (*13*) (Fig. 7E and F).

## Discussion

Amyloid fibrils assembled from unrelated proteins converge on the same cross-β architecture; however, which residues that architecture can accept is not readable from any one sequence. We approached the question by measuring one protein completely. DMS of every amino acid substitution in α-synuclein gave a quantitative map of the determinants of aggregation that extends well beyond the familial Parkinson’s disease mutations. On that map, the substitutions that suppress aggregation converge on glycines of the fibril core. Human variation samples at most a few substitutions per position, so intolerance that is uniform across a position can be identified only by measuring all of them. Within the human single-nucleotide-variant screen, G68E ranked as the strongest suppressor. We then turned to the structures, asking what makes a Cβ unacceptable at those positions.

Structural modeling explained this suppression as a packing constraint rather than the loss of a specific structural feature: Gly68 and Gly73 sit where the lattice leaves no room for the Cβ atom that every other amino acid carries. At Gly68 that room cannot be created: in five of the six structures the adjacent layer sets the limit and expanding the stack breaks the inter-subunit hydrogen bonds before a Cβ fits. In 6H6B the limiting atom belongs to the subunit itself, where no axial expansion can help, but repacking finds the substitution destabilizing there as well (Fig. 4F). The constraint was reproduced in all six structures examined, indicating that it reflects a geometric property of the ordered core rather than a feature of a single fold (Fig. 4C and Fig. S3A).

The protection G68E confers is confined to the molecule that carries it. G68E suppressed aggregation on all familial-mutant backgrounds tested, yet when co-expressed with WT α-synuclein it did not lower the WT aggregate burden; it therefore acts in cis rather than as a dominant, trans-acting inhibitor. At the same time, G68E α-synuclein retained normal localization to the soma, neurites and synaptic sites in iPSC-derived neurons, consistent with Gly68 lying on the cytoplasmic face away from the membrane-binding interface (Fig. S6B to D). G68E thus removes the molecule’s capacity to fibrillize while leaving its physiological behavior intact, a separation of aggregation from function that many aggregation-modifying substitutions do not achieve.

The same substitution occurs outside humans, and its effect there had been noted before: α-synuclein from elephant, bowhead whale and pig fibrillates poorly, and of the three substitutions these species share relative to the human protein, A53T, G68E and V95G (*22*). Several mammals carry G68E or V95G while retaining the aggregation-prone ancestral Thr53 (*20*), and replacing those residues with the human ones restores aggregation (Fig. 5A to D). These species are therefore protected not by lacking the aggregation-prone residue, which they still carry, but by the substitution at position 68 or 95.

The constraint is not peculiar to α-synuclein. It holds in 51 of the 53 amyloid systems we surveyed, in proteins that share no sequence, no native fold and no function. The two exceptions are the systems whose core offers a single glycine, so they are small samples rather than counterexamples; every system with more than one contains at least one blocked glycine. The common feature among the 51 systems is the repeat itself: the backbone hydrogen bonding that defines a cross-β core fixes the spacing between layers near 4.8 Å (*1*), so which positions the lattice closes is set by geometry rather than by sequence. The constraint is therefore imposed by assembly rather than carried by the monomer. Buried glycines of folded proteins are blocked as often (Fig. S8A and B); what differs is that the blocking atom shifts from a movable same-layer side chain to the backbone of the adjacent layer, which cannot move without dismantling the structure. Applied to a second amyloid, the same positional logic identifies Gly29 in the amyloid-β core (*5*), where our cell measurements agree with the published nucleation scan (*25*): what the position refuses is the Cβ, almost irrespective of what the side chain carries beyond it. Because the measurement needs only coordinates, any newly determined fibril structure can be screened before an experiment is designed, though it gives no ranking within a fold: the tightest glycine of one structure need not be the one substitutions avoid.

Two earlier deep mutational scans of α-synuclein used different readouts and so answer different questions. Chlebowicz et al. quantified seeded aggregation in a HEK293T FRET biosensor, finding inhibitory hotspots within residues 65 to 78 and a modulating role for the N-terminal region (*19*); Newberry et al. used yeast growth, showing that toxicity there arises from a membrane-bound amphipathic helix rather than from amyloid, and is relieved by mutations that destabilize the helix (*24*). Those readouts report on seeding and on membrane-associated toxicity. Our saponin-based reporter reads fibril content directly, which is what resolves the core-glycine constraint and links the strongest protective variant to its structural basis, its evolutionary recurrence, its behavior in neurons and its effect in a mouse seeding model.

This study has several limitations. First, the DMS assay relies on PFF□induced aggregation in HEK293T cells, which may not fully capture the complexity of α□synuclein behavior in neurons or in vivo. Second, the structural survey covers the deposited fibril record, not the full diversity of polymorphs in human disease: entries are retrieved by the words amyloid and fibril in the entry text, and structures whose coordinates contain fewer than three in-register layers are excluded, so some fibrils are represented only by their multi-layer depositions. Third, although G68E shows strong protective effects in mice, its impact in human neural circuits and disease progression remains to be determined.

Within these limitations, the work defines the mutational architecture of α-synuclein aggregation and grounds the most suppressive variant of the human single-nucleotide-variant screen in a specific, physically interpretable constraint. Because G68E protects only in cis, a therapeutic strategy built on it would have to alter the endogenous allele, through gene editing or allele replacement, rather than supply the variant in trans. More generally, the core-glycine packing constraint marks a defined structural liability of the fibril that small molecules or peptide mimetics might exploit, and the variant-effect map provides a resource for stratifying risk and identifying candidate protective alleles across synucleinopathies.

## Materials and Methods

### Cell culture

HEK293 cells and HEK293T cells (Clontech, Mountain View, CA) were cultured and maintained at 37□with 5% CO_2_ in Dulbecco’s modified Eagle’s medium (DMEM, WAKO, Osaka, Japan), supplemented with 10% fetal bovine serum (FBS, Thermo Fisher Scientific, Waltham, MA) and penicillin-streptomycin (100 U/ml, Thermo Fisher Scientific). All cell lines were routinely tested and confirmed to be negative for mycoplasma contamination.

### Animals

Animal experiments were performed in accordance with international guidelines for the care and use of experimental animals. Mice were maintained under a 12-h light-dark cycle (lights on from 08:00 to 20:00 h) with free access to water and food. All animal experiments, including production and maintenance protocols, were reviewed and approved by the Animal Care and Use Committee of the Kyoto Prefectural University of Medicine (M2024-517). Mice were anesthetized with a midazolam/medetomidine/butorphanol cocktail (4, 0.3 and 5 mg/kg, respectively) during surgery.

### Plasmids

SNCA-GFP, SNCA-GFP-IRES-miRFP670 genes were cloned into the *Xba*I-*Sal*I sites of the pLenti vector. A series of mutant SNCA constructs harboring specific amino acid substitutions was generated from pEGFP-SNCA(*41*) using oligonucleotide-directed PCR mutagenesis. SNCA-IRES-GFP genes were cloned into the *BamH*I-*EcoR*I sites of the pAAV EF1 vector.

### Lentivirus production

HEK293T cells (Clontech) were seeded in 10 cm dishes and transfected at approximately 70% confluence with 9 µg of the plasmid library, 6 µg of psPAX2 (Addgene #12260), and 3 µg of pMD2.G (Addgene #12259) using FuGENE HD (Promega, Madison, WI), according to the manufacturer’s instructions. Culture supernatants were collected after 48 h, centrifuged at 3,000 rpm for 10 min at 4□, and filtered through a 0.45 µm low protein-binding filter (SFCA). Viral supernatants were stored at –80□until use.

### AAV packaging

The AAV vector was used to overexpress WT or G68E α□synuclein in the brain of SNCA knockout mice. pAA EF1 SNCA-IRES-GFP was packaged by VectorBuilder Inc. (Chicago, IL, USA).

### AAV injection in the mouse brain and immunohistochemistry

A total of 2 μL of AAV SNCA-IRES-GFP WT or G68E (1.5 × 10¹³ GC/ml) was injected into the bilateral striatum of B6;129X1-*Snca*^tm1Rosl^ /J mice (JAX stock #003692)(*42*) using a 26G injection needle attached to a stereotaxic syringe pump (Legato 130; KD Scientific, Holliston, MA). Expression of α-synuclein from the injected AAV was confirmed by immunostaining using an anti–α-synuclein antibody (#sc-7011-R, Santa Cruz Biotechnology, Dallas, TX) (Fig. S7). One week after the injection, 2 μL of α-synuclein PFF (2 μg/μL) was additionally injected into the striatum on one side(*43*). Three months after PFF injection, mice were perfused and fixed with 4% paraformaldehyde, and brain samples were collected. Brain sections (50 μm thick) were prepared using a vibratome (PRO7; Dosaka EM, Kyoto, Japan). Blocking and permeabilization were performed with 2.5% normal goat serum (NGS) and 0.1% Triton X-100 for 1 h at room temperature. Brain sections were incubated with anti-phospho-α-synuclein (#ab59264, Abcam, Cambridge, UK) and anti-p62 (#ab56416, Abcam) antibodies for 2 days at 4 °C, followed by incubation with secondary antibodies (anti-rabbit IgG Alexa Fluor Plus 488 and anti-mouse IgG Alexa Fluor 350, #A-11045, Thermo Fisher Scientific) after washing with PBS. Samples were mounted with FluorSave Reagent (Calbiochem, San Diego, CA). mouImages were acquired using a Zeiss LSM 900 laser scanning confocal microscope equipped with 10×/0.45 or 40×/0.95 objectives (Carl Zeiss AG, Oberkochen, Germany). Phospho-α-synuclein–positive signals in each mouse (*n* = 5) were quantified from stained images using the Analyze Particles function in ImageJ software (NIH). Signals larger than 50 pixels were defined as aggregates and counted.

### Introduction of α-synuclein fibrils and aggregation assay

HEK293T cells expressing α-synuclein-GFP were harvested, centrifuged, suspended with FBS-free DMEM to a concentration of 1.0 × 10^6^/ml, and seeded in 12-well plate or 96-well plates at 1 ml or 100 µl per well. In case of 96-well plate, a 40 µl solution of FBS-free DMEM containing 2 µg of sonicated α-synuclein PFF and 0.8 µl of DMRIE-C (10459014, Thermo Fisher Scientific) was then added to each well. After 6 hours, 140 µl of DMEM with 20% FBS was added to each cell. Two days after incubation, α-synuclein-GFP aggregates were assessed by flow cytometry (Attune, Thermo Fisher Scientific) or InCellAnalyzer2200 (GE Healthcare, Chicago, IL). For flow cytometry analysis, cells were resuspended and treated with a final concentration of 0.1% saponin solution in PBS for 20 minutes prior to analysis. For InCellAnalyzer analysis, cells were stained with Hoechst33342 dye for the measurement of aggregates. All live cell data was acquired using the DAPI and FITC filter set. The images obtained were analyzed using the InCellAnalyzer workstation. Aggregates and nuclei were counted automatically, and data was represented as granule count/cell.

### Sarkosyl fractionation and immunoblot analysis of α-synuclein

HEK293 cells were plated at 8 × 10^5^ cells per well in 6 well plates and cultured overnight. Cells were transfected with plasmids encoding WT, the Δ73–83 deletion mutant (D; non aggregating), or the G68E variant using 1□µg DNA per construct (WT: 1□µg; D: 1□µg; G68E: 1□µg; WT+D: 1□µg WT + 1□µg D; WT+G68E: 1□µg WT + 1□µg G68E; WT+WT: 1□µg WT + 1□µg WT). After 3□h, WT α□synuclein PFF (0.12 µg per well) were added as seeds, and cells were incubated for 3□days. Cells were harvested in 600□µL of A68 buffer containing 1% sarkosyl (10□mM Tris□HCl pH□7.5, 10% sucrose, 0.8□M NaCl, 1□mM EGTA). Lysates were sonicated and subjected to ultracentrifugation at 100,000□×□g for 20□min. The sarkosyl□soluble supernatant (sup) and sarkosyl insoluble pellet (ppt) fractions were collected separately. Protein samples were analyzed by immunoblotting using antibodies against the α□synuclein C□terminus (epitope 131–140: anti a-synuclein 131-140, TIP-SN-P09, Cosmo Bio, Tokyo, Japan) and phosphorylated Ser129 (anti-pSyn#64, 015-25191, FUJIFILM Wako).

### DMS Library construction, FACS, and Illumina sequencing

The SNCA coding sequence was divided into two segments, and saturation mutagenesis was independently introduced into each segment. Pooled oligonucleotides containing degenerate NNK codons were synthesized by Integrated DNA Technologies. Synthesized oligonucleotides were extended by overlap PCR and cloned into the *Xba*I-*EcoR*I site of pLenti CMV IRES-miRFP670, modified from the plasmid (addgene #17448) using the NEBuilder HiFi DNA Assembly Kit (New England Biolabs, Ipswich, MA). An 80-µl assembly reaction was performed using 10 µg of gel-purified inserts and 5 µg of vector. The assembled products were purified using a Gel and PCR Clean-up kit (Takara Bio, Kusatsu, Japan) and eluted in 20 µl of nuclease-free water. Subsequently, 200 µg (approximately 4 µl) of the purified assembly reaction was transformed into 50 µl of electrocompetent cells (Lucigen, Middleton, WI) and expanded according to the manufacturer’s protocol using an ECM 399 system (BTX, Holliston, MA) with 1500 V electroporation. A 1,000-fold dilution of the full transformation was plated to estimate the library complexity. To prevent recombination between libraries, the two sub-libraries were packaged into lentiviral particles separately and individually transduced into HEK293T cells with a multiplicity of infection (MOI) of less than 0.2 to ensure that no more than one SNCA mutant was expressed per cell. At 48 h post-transduction, miRFP670-positive cells were enriched by fluorescence-activated cell sorting (FACS) using an MA900 cell sorter (Sony, Tokyo, Japan). The two sub-libraries were then pooled and cultured for an additional 5 days. For an aggregation analysis, library cells were suspended with FBS-free DMEM to a concentration of 1.0 × 10^6^/ml, and seeded in 10 cm dish at 10 ml. A 1 ml solution of FBS-free DMEM containing 50 µg of sonicated PFF and 20 µl of DMRIE-C (10459014, Thermo Fisher Scientific) was then added. After 6 hours, 5.5 ml of DMEM with 30% FBS was added. Two days after incubation, cells were sorted according to the signal of GFP with 0.1% saponin pretreatment using a Sony SH800 cell sorter. For an abundance analysis, library cells were resuspended in the complete medium and sorted using a Sony SH800 cell sorter. For genomic DNA extraction, sorted cells were incubated overnight at 55□°C in 6□mL NK Lysis Buffer (50□mM Tris, 50□mM EDTA, 1% SDS, pH□8) containing 30□µL Proteinase K (20□mg/mL). The following day, 30□µL RNase A (10□mg/mL) was added, mixed by inversion, and incubated at 37□°C for 30□min. After cooling on ice, 2□mL of pre□chilled 7.5□M ammonium acetate was added to precipitate proteins. Samples were vortexed and centrifuged at ≥4,000□×□g for 10□min, and the clarified supernatant was transferred to a fresh tube. Genomic DNA was precipitated with 6□mL 100% isopropanol, mixed by inversion, and centrifuged at ≥4,000□×□g for 10□min. Pellets were washed with 6□mL freshly prepared 70% ethanol, briefly centrifuged, and residual ethanol was removed. DNA pellets were air dried for 10–30 min until translucent, then resuspended in 500□µL 1×□TE buffer. Samples were incubated at 65□°C for 1□h and at room temperature overnight to fully dissolve the DNA. Concentrations were measured using a NanoDrop spectrophotometer(*44*). Each library was amplified using sub-library-specific primers, followed by a second round of PCR to add Illumina flow-cell adapters and sample-specific barcodes. The resulting amplicons were sequenced on an Illumina NovaSeq 6000 platform using a 150 bp paired-end protocol at the Department of Infection Metagenomics, Research Institute for Microbial Diseases, Osaka University. Sequencing data were analyzed by comparing read counts between GFP-high and GFP-low populations, with normalization performed relative to the WT sequence read count.

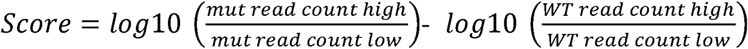

Aggregation and abundance scores were computed as described above for each of the 2,641 missense substitutions, for the nonsense substitutions and for the synonymous WT controls. Each score is the mean of two independent replicate experiments. The standard error of that mean was estimated from the replicate difference as |rep1 − rep2| / 1.128 / √2, the unbiased estimator of the standard deviation from a two-observation range, giving a median standard error of 0.044 for aggregation and 0.079 for abundance. A variant was called significant when |score| / standard error exceeded the two-sided normal critical value at a Benjamini–Hochberg false-discovery rate of 5% (Benjamini and Hochberg, 1995; no PubMed record) across all 2,641 substitutions; 897 substitutions were significantly suppressive and 792 significantly aggregation-promoting on this criterion. No read-depth filter was applied, the lowest depth over all variants being 181 reads for aggregation and 138 for abundance (medians 5,679 and 5,369). The nonsense substitutions, which report the absence of protein, define the internal floor of each assay (aggregation −0.83 ± 0.17, abundance −1.04 ± 0.37).

### Generation of *SNCA* G68E iPSCs

To generate iPSCs harboring the *SNCA* G68E mutation, CRISPR-Cas9–mediated homology-directed repair (HDR) was performed using the Alt-R™ CRISPR-Cas9 system (Integrated DNA Technologies, IDT, Coralville, IA). A crRNA targeting the *SNCA* locus was designed and synthesized by IDT. The crRNA was annealed with Alt-R™ CRISPR-Cas9 tracrRNA to form guide RNA duplexes according to the manufacturer’s instructions. For fluorescence-based assessment of transfection efficiency, Alt-R™ CRISPR-Cas9 tracrRNA, ATTO™ 550 was used in parallel experiments. Ribonucleoprotein (RNP) complexes were assembled by combining recombinant Alt-R™ S.p. Cas9-GFP V3 (#10008100, IDT) with the crRNA:tracrRNA duplexes. A single-stranded HDR donor oligonucleotide, containing the *SNCA* G68E mutation and flanking homology arms (5’-ATTTGTACCAGTGGCTGAGAAGACCAAAGAGCAAGTGACAAATGTTGGAGAAGCAG TGGTGACGGGTGTGACAGCAGTAGCCCAGAAGACAGTGGAGGGA-3’), was co-delivered to enable precise genome editing via HDR. Electroporation Enhancer and HDR Enhancer (IDT) were included to improve editing efficiency in accordance with the manufacturer’s instruction.

Human iPSCs generated from healthy donor, hc2EL5 were dissociated into single cells and 5 × 10^5^ cells were electroporated with the RNP complexes and HDR donor using 4D-Nucleofector system (Lonza, Basel, Switzerland) with the P3 Primary Cell 4D-Nucleofector X Kit (Lonza) according to the manufacturer’s instructions. After 48h electroporation, cells were subjected to single-cell sorting by fluorescence-activated cell sorting (FACS) based on GFP and RFP double-positive selection and maintained with StemFit AK02N medium (Ajinomoto, Tokyo, Japan) on laminin (iMatrix-511; Takara Bio Inc., Kusatsu, Japan)-coated culture plates at 37◦C in a standard 5 % CO2 incubator for 7 days. For genomic DNA extraction, iPSC pellets were resuspended in PBS and incubated with 1 mg/mL Proteinase K at 55°C for 2 hours or overnight. Following heat inactivation at 95°C for 10 minutes, the targeted region was amplified by PCR. Homozygous SNCA G68E mutation clones were confirmed by Sanger sequencing.

### Conversion of iPSCs into cortical neurons

A tetracycline-inducible human neurogenin 2 (*NGN2*) construct was transduced into iPSCs using a *piggyBac* vector(*45, 46*). The iPSC colonies incorporating the construct were selected with neomycin, G418 disulfate (Nacalai Tesque, Kyoto, Japan). For differentiation into cortical neurons, iPSCs were dissociated into single cells and replated onto laminin (iMatrix-511)-coated 24-well plates at a density of 50,000 cells/well. The differentiation medium consisted of a 1:1 ratio of DMEM/F12 (Thermo Fisher Scientific) and Neurobasal (Thermo Fisher Scientific), 1% N2 supplement (Thermo Fisher Scientific), 2% B27 supplement minus vitamin A (Thermo Fisher Scientific), 1× L-glutamine (Thermo Fisher Scientific), 1 × Penicillin–Streptomycin (Thermo Fisher Scientific), 10 ng/ml brain-derived neurotrophic factor (BDNF, R&D Systems, Minneapolis, MM), 10 ng/ml glial cell-derived neurotrophic factor (GDNF, R&D Systems) and 10 ng/ml neurotrophin-3 (NT-3, R&D Systems) with 1 μg/mL doxycycline hydrochloride (Wako Pure Chemicals Industries, Osaka, Japan). The medium was also supplemented with 10 μM Y-27632 (Nacalai Tesque) during the initial plating. The medium was exchanged twice per week.

### Immunocytochemistry and image acquisition

iPSC-derived neurons at DIV28 were fixed with 4% paraformaldehyde and washed three times with PBS. For blocking and permeabilization, cells were incubated in blocking solution (2.5% normal goat serum and 0.1% Triton X-100 in PBS) for 30 min at room temperature. Primary antibodies were diluted in blocking solution (anti-α-synuclein, #sc-7011-R, Santa Cruz Biotechnology, Dallas, TX; anti-Synapsin-1, #106011, Synaptic Systems GmbH, Göttingen, Germany; anti-MAP2, #ab5392, Abcam, Cambridge, UK; anti-TUBB, #TUJ-0020, Aves Labs, Tigard, OR). Cells were incubated with primary antibodies for 18 h at room temperature. After washing with PBS, cells were incubated with secondary antibodies (anti-rabbit IgG Alexa Fluor Plus 488; anti-mouse IgG Alexa Fluor Plus 594, Thermo Fisher Scientific, Waltham, MA; anti-chicken IgY DyLight 405, Rockland Immunochemicals, Limerick, PA) for 3 h at room temperature. Samples were mounted with FluorSave Reagent (Calbiochem, San Diego, CA). Images were acquired using a Zeiss LSM 900 laser scanning confocal microscope equipped with a 63×/1.4 NA oil immersion objective (Carl Zeiss AG, Oberkochen, Germany).

### Human single-nucleotide variants and population frequencies

These 88 SNPs were extracted from gnomAD, dbSNP, and ClinVar. Allele frequencies were taken from gnomAD (*47*), using the global (ALL) frequency of the exome call set where available and of the genome call set otherwise: 47 of the 88 variants are reported in gnomAD exomes, 23 in gnomAD genomes and 18 are absent from both. Reported frequencies span 6.84 × 10^-7^ to 2.558 × 10^-4^.

### Fibril structures and construction of the layer stack

α Synuclein analyses used six fibril structures: the three multiple system atrophy filaments 6XYO, 6XYP and 6XYQ, the Lewy fold 8A9L, and the in vitro polymorphs 6SSX and 6H6B. Amyloid-β analyses used 7Q4B and 7Q4M (Alzheimer brain), 5OQV (Aβ42 in vitro) and 6SHS (Alzheimer brain, Aβ40) by cryo-electron microscopy, and 2NAO and 5KK3 determined by NMR; for the NMR entries the first model of the deposited ensemble was used. For each structure, waters, ligands and hydrogens were removed and heavy atoms only were retained; hydrogens were present in the deposited 6H6B coordinates and were removed. Three in-register subunits of one protofilament were selected and renamed A, B and C in stacking order, and all measurements were made on the middle subunit B, which alone has a neighbor on both faces. Subunits were taken from the deposited coordinates except for 8A9L, whose model contains a single layer and whose stack was generated by applying the deposited helical rise (4.77 Å) and twist (0.85°) about the filament axis. The axial rise per subunit is a property of the lattice and was measured as the subunit-to-subunit translation; it is 4.73 Å in 6XYO.

### Free radius at the modeled Cβ position

For every residue of the scored subunit, the position a Cβ atom would occupy was constructed from that residue’s own backbone N, CA and C atoms by the standard tetrahedral relation, so that the measurement is defined identically at glycine and at non-glycine positions. The free radius is the radius of the largest sphere centered on that point that overlaps no neighboring van der Waals surface, that is the minimum over all other atoms of the interatomic distance less the van der Waals radius of that atom. Radii were 1.70 Å for carbon, 1.55 Å for nitrogen, 1.52 Å for oxygen, 1.80 Å for sulfur and phosphorus and 1.90 Å for selenium. Three classes of atom were excluded from the minimum: the residue’s own atoms; the atoms of the covalently bonded neighbors i − 1 and i + 1 of the same chain, which approach any Cβ position by bonded geometry alone; and atoms within 2.4 Å of one of the residue’s own atoms, which removes disulfide partners. A position whose free radius is below 1.70 Å, the van der Waals radius of carbon, cannot accept a Cβ atom without displacing the blocking atom. Two checks bound the construction: over the structures analyzed here the modeled position differs from the observed Cβ of non-glycine residues by a median of 0.045 Å, and on 6XYO the measurement returns 0.44 Å at Gly68 with ND2 of Asn65 in the adjacent subunit as the limiting atom, and 1.15 Å at Gly67 where the limiting atom belongs to the same subunit.

### Burial and assignment of the blocking atom

Solvent accessibility was computed with an implementation of the Shrake–Rupley algorithm (*48*) written for this work, using a 1.4 Å probe and 200 points per atom, in the context of the complete deposited model, and normalized to the theoretical maximum accessible surface area of the residue type (*49*). Residues with a relative accessibility below 0.25 were taken to be buried, and all glycine analyses are restricted to buried positions. The origin of a blockage was assigned geometrically rather than by chain identity: the fibril axis was taken as the subunit-to-subunit translation, and the limiting atom was assigned to an adjacent layer when it belongs to a different subunit, or when, within the same chain, it is displaced along the axis by more than half a rise. The second clause is required for β solenoid folds, in which consecutive layers belong to one chain and a chain-identity rule would misclassify inter-layer contacts as intra-chain. Each position is additionally classified by whether it is blocked from an adjacent layer, from within its own layer, from both, or from neither.

### Rigid axial expansion and the inter-subunit hydrogen-bond network

To test whether the axial block can be relieved by making room, each three-subunit stack was expanded rigidly along the fibril axis, the two outer subunits being translated away from the middle one in 0.01 Å steps without changing any internal coordinate, and the free radius at the modeled Cβ position of Gly68 was recomputed at each step. In parallel, inter-subunit backbone hydrogen bonds were enumerated in the deposited stack as backbone amide N to backbone carbonyl O pairs of different subunits within 3.2 Å (n = 57 to 137 per structure), and a bond was counted as broken when expansion carried that distance beyond 3.5 Å. A carbon first fits at Gly68 at rises of 4.93 to 6.57 Å depending on the structure, against observed rises of 4.7 to 4.8 Å. In the three brain-derived filaments and in 6SSX the network is entirely broken before that point, the first bond breaking at 4.99 to 5.11 Å where the free radius is still 0.52 to 0.75 Å; in 8A9L 77% of the bonds are broken; in 6H6B, the lowest-resolution structure of the set, a carbon fits at the observed rise. Molecular images in Fig. 4D and Fig. S3B were rendered in PyMOL 3.1.0 with the camera set along the filament axis of the three-subunit stack; labels, leader lines and panel annotations were added in matplotlib.

### Backbone conformation analysis

As a negative control for the steric criterion, the backbone conformation of each core glycine was compared with the conformational preferences of the residue types that could replace it. Reference φ/ψ distributions were compiled from 287,671 residues of 1,400 X-ray structures at 1.6 Å resolution or better, excluding glycine, proline and the residues preceding proline. Contours enclosing 98% and 99.95% of that set are shown in Fig. S3D, together with the region in which all 18 non-proline, non-glycine types are observed within 30° and the corresponding region for proline. For each glycine of each structure, the number of reference residues of each type lying within 30° of its observed conformation was counted.

### Repacking calculations

The energetic cost of introducing a Cβ atom was estimated with PyRosetta (*50*). For each three-subunit stack, the residue of interest was mutated in all three subunits, the stack was relaxed with FastRelax (5 repeats, ref2015 score function, with the -ex1 -ex2 rotamer expansion), and the relaxed pose was analyzed with InterfaceAnalyzerMover (*17*), subunit B being scored against subunits A and C. Fifty independent models were generated per state with independent random seeds, and the reported difference is between the means of the 50 mutant and 50 WT models, with the 95% confidence interval of that difference. Absolute total scores are not compared between folds, because the number of modeled residues per subunit differs; only the mutant-minus-WT difference within a fold is interpreted. The same protocol was used for G68E in the six α□synuclein structures (Fig. 4F) and for G29E in the six amyloid-β folds (Fig. 7D and Fig. S8D)

### Cross-amyloid survey: retrieval and selection of fibril structures

Structures were selected by a query against the Protein Data Bank followed by a geometric test on the deposited coordinates, so that membership depends neither on deposition annotations nor on which systems we judged representative. The RCSB Protein Data Bank Search API was queried (data frozen 29 September 2026) for entries whose full text contains “amyloid” or “fibril” and whose experimental method is electron microscopy, solid-state NMR or solution NMR, returning 3,008 entries. Structure-type annotations were deliberately not used to narrow the retrieval, because they are inconsistent for fibril structures. An entry was then retained when it met all of the following: resolution 4.5 Å or better for electron microscopy, NMR entries being exempt; at most three distinct protein entities and at most 6,000 modeled polymer residues; the deposited coordinates contain an in-register cross-β ladder, defined as three subunits of the same polymer whose residue-by-residue CA displacements form a pure translation (median absolute deviation below 0.8 Å on each component over at least 8 shared residues), with the two outer subunits on opposite sides of the middle one and a translation of 4.2 to 5.2 Å per layer; and at least one glycine of the scored subunit buried as defined above. Where one subunit spans several rungs, as in β solenoid folds, the displacement was divided by the number of rungs before the ladder test. The first two criteria left 2,366 entries, of which 2,363 were retrieved and 2,350 parsed; the ladder criterion was met by 849 and the glycine criterion by 811 of those. Because the ladder must be present in the deposited model, structures containing one or two layers are excluded rather than completed by applying the reported helical symmetry, so every free radius reported here is measured on deposited atoms; HET-s and CsgA are represented in the set only by their multi-layer depositions.

The 811 structures carry 70 UniProt entries, corresponding to 53 proteins once entries of the same protein from different species, isoforms, cleaved fragments and point-mutant constructs are merged, together with patient-derived immunoglobulin light chains that carry no UniProt reference; counts of systems in the text are protein-level. Because one fold is often deposited many times, entries were grouped by UniProt entry and, within an entry, by the set of buried core glycine positions together with their inter-layer free radii; entries agreeing on the position set and within 0.2 Å at every position were treated as the same fold and the higher-resolution entry retained, leaving 687 non-redundant folds. The result does not depend on the resolution cutoff: restricting the non-redundant set to entries at 4.0, 3.5, 3.0 or 2.5 Å or better, with NMR entries retained, gives 70.0%, 70.1%, 71.7% and 73.6% of buried core glycines blocked from an adjacent layer, against 70.1% for the full set. Results are reported for both sets and agree: 70.9% of all 5,363 buried core glycine positions and 70.1% of the 4,697 positions of the non-redundant subset are blocked from an adjacent layer. Where a structure contains several distinct protofilament sequences, one ladder per sequence was scored, and a position observed in more than one ladder of the same structure contributes once. The positive control for the measurement is the buried non-glycine positions of the same cores, at which a Cβ atom is in fact present.

### Natively folded control set

The fold control is a set of 21 natively folded structures, comprising the native folds of fibril-forming proteins in the survey and globular proteins of unrelated fold, scored one chain at a time with every deposited chain present as environment. It contains 92 buried glycines and was not re-selected when the fibril set was re-selected.

### Paired native-to-fibril comparison

The native side of the paired comparison was selected by the same logic as the fibril side. For every UniProt entry in the fibril set, all Protein Data Bank entries carrying that entry were retrieved (3,281 entries over 69 accessions) and those that are absent from the fibril set, contain a single protein entity and no nucleic acid, have at most 600 modeled polymer residues, were solved by X-ray diffraction, electron microscopy or solution NMR, and reach 3.0 Å or better where a resolution applies were kept; 1,740 entries passed, and the three highest-resolution entries per accession were scored (100 structures). Two structural criteria then decide. A candidate must be a folded monomer, defined as at least 25% of the residues of the scored chain being buried when that chain is scored alone; this removes isolated amyloidogenic peptides and extended conformers that a metadata filter cannot distinguish from a native fold, and 49 of the 100 candidates fail it, the six- and seven-residue steric-zipper crystals scoring 0%. A candidate must also cover the fibril core: native and fibril structures were aligned by the constant residue-numbering offset that maximizes residue-type agreement over shared positions, requiring at least 95% agreement over at least 10 positions, and a glycine enters the comparison only if it is a glycine in the native structure and a buried core glycine in the fibril. For each accession the candidate covering the most such glycines was kept, with resolution as tie-break. Because a protein contributes many fibril folds, the fibril value of a glycine is the median over the non-redundant folds of that accession, so each glycine enters the test once. The comparison covers 104 glycines of 13 proteins over 95 fibril folds, and native and fibril values were compared by the Wilcoxon signed-rank test.

### Positioning of the membrane-bound helix in an implicit bilayer

Residues 61 to 84 of chain A of the micelle-bound α-synuclein structure (PDB 1XQ8, model 1) were positioned in an implicit DOPC bilayer with PPM 3.0, using a single membrane and no topology restraint. The full chain was also submitted but could not be placed as a rigid body: its transfer energy was weaker than that of the 24-residue segment (−6.0 against −9.3 kcal mol ¹) and only five residues reached the hydrocarbon core, two of them polar, because the two helices of the micelle-bound structure present hydrophobic faces 85° apart and cannot be buried by a single plane. Depths were measured from the hydrocarbon-core boundary defined by the DUM pseudo-atoms of the output coordinates; this reproduced the embedded-residue list returned by the server for every residue except Val77, which lies 0.03 Å from the plane. For Fig. S6C, side-chain directions were projected onto the plane normal to the helix axis, with the membrane normal as the reference direction; for the glycines the Cβ position was built from the backbone, a construction that reproduces the observed Cβ directions of the other residues to within 3°.

### Sequence logo representation and cladogram of α synuclein conservation

Vertebrate α-synuclein sequences were retrieved from the aLeaves server(*51*), aligned using MAFFT(*52*), and visualized with the WebLogo 3 server(*53*). The 53rd, 68th and 95th amino acid residues of each taxon are shown along with the generally accepted phylogenetic relationships among placental mammals and among reptiles and birds. Species silhouettes were obtained from Creazilla: Natasha Sinegina (CC BY 4.0; URLs listed below) and public domain sources. https://creazilla.com/media/silhouette/294 https://creazilla.com/media/silhouette/2421 https://creazilla.com/media/silhouette/2526 (cropped) https://creazilla.com/media/silhouette/1366 https://creazilla.com/media/silhouette/2459 https://creazilla.com/media/silhouette/2963 https://creazilla.com/media/silhouette/457

### Phylogenetic comparative analysis

Orthologs of human SNCA were retrieved from Ensembl Compara (release 116) and the residue aligned to human position 68 was read from each pairwise protein alignment. Maximum longevity and adult body mass were taken from AnAge (build 15); species lacking either value were excluded, leaving 69 placental mammals (33 Glu68, 36 Gly68). The Ensembl Compara vertebrate species tree was pruned to these taxa, made ultrametric (root-to-tip coefficient of variation before adjustment, 1.5%) and scaled to a 105-Myr placental root. Phylogenetic signal was estimated as Pagel’s λ by maximum likelihood (λ = 1.00 for both log_10_ maximum lifespan and log_10_ adult body mass; likelihood-ratio P = 2.2 × 10^-11^ and 8.6 × 10^-10^), and phylogenetic clustering of the binary residue-68 state as Fritz and Purvis’ D (D = −9.36; P = 0.001 against a random distribution, P = 1.00 against a Brownian threshold model). The association between residue-68 state and longevity was tested by PGLS with λ estimated by maximum likelihood, with and without log_10_ adult body mass as a covariate, and by phylogenetic ANOVA (10,000 Brownian-motion simulations on the tree). Ancestral states were reconstructed by maximum parsimony and, as a check, under a two-state Mk model fitted by maximum likelihood, both on the 69-taxon placental tree and on the full Ensembl Compara vertebrate tree pruned to the 169 ortholog species present in it. Analyses used DendroPy. The code is provided as a Source Data file.

### Quantification and statistical analysis

Data are presented as individual data points overlaid on mean ± SEM or box-and-whisker plots. The sample size (n) refers to biological replicates as indicated in each legend. Cell and animal experiments were analyzed with the unpaired two-tailed Mann-Whitney test and two-way repeated-measures ANOVA followed by Šídák’s multiple-comparison test, in GraphPad Prism 10 (GraphPad Software, LLC, San Diego, CA). For the structural analyses, distributions of free radius were compared with the two-sided Mann-Whitney U test and paired native-to-fibril values with the Wilcoxon signed-rank test; variant scores were tested as described above and corrected by the Benjamini-Hochberg procedure (Benjamini and Hochberg, 1995; no PubMed record) at a false-discovery rate of 5%. Correlations are Spearman coefficients. These analyses, and all structural measurements, were performed in Python with gemmi, NumPy, SciPy, pandas and Matplotlib, and repacking calculations with PyRosetta.

## Data availability

Source data are provided with this paper. The Source Data file gives, on one sheet per panel, the measurements plotted in Fig. 1D, Fig. 3A to E, Fig. 4C, E and F, Fig. 5A and B, Fig. 7A to D, Fig. S3C to E, Fig. S5A to C, Fig. S6 and Fig. S8A to D, together with an index sheet that lists every panel and the table underlying it and identifies the panels that carry no plotted data. The scripts implementing the measurement, the selection and the phylogenetic comparative analysis are provided as source data. Fibril and native structures were taken from the Protein Data Bank under the accessions listed in the Methods, population allele frequencies from gnomAD v4, orthologs from Ensembl Compara release 116 and life-history data from AnAge build 15.

## Acknowledgments

This research was supported by a JSPS Grant-in-Aid Scientific Research Grant 20K21585 (A.H.), 24K10494 (A.H.), and 26K02063 (K.K.), AMED under Grant Number JP24wm0625501 (A.H., H.I.), JP23bm1323001 (H.I.), JP23bm1223013 (H.I.), and JP23dk0207066 (H.I.), and Takeda Science Foundation (A.H.). Experimental schemes were partially illustrated by BioRender. During the preparation of this manuscript, the authors used Paperpal to polish the language. After using this tool, the authors reviewed and edited the content as needed and took full responsibility for the content of the publication.

## Author contributions

A.H. conceptualized the study. S.H., N.H., M.O., and A.H. conducted the DMS and subsequent *in vitro* assays. T.N., K.K., Y.K., and A.H. contributed to data analysis and mathematical modeling. Y.W., R.S., and K.T. conducted mouse experiments. T.N. evaluated the aggregation property with western blot. T.O., K.I., T.K., K.T. performed iPSC differentiation to neuron. A.H. wrote the manuscript with the help of all the authors. M.H., H.I., S.M., and A.H. supervised the study. All authors reviewed and approved the final manuscript.

---

**Fig. S1.**
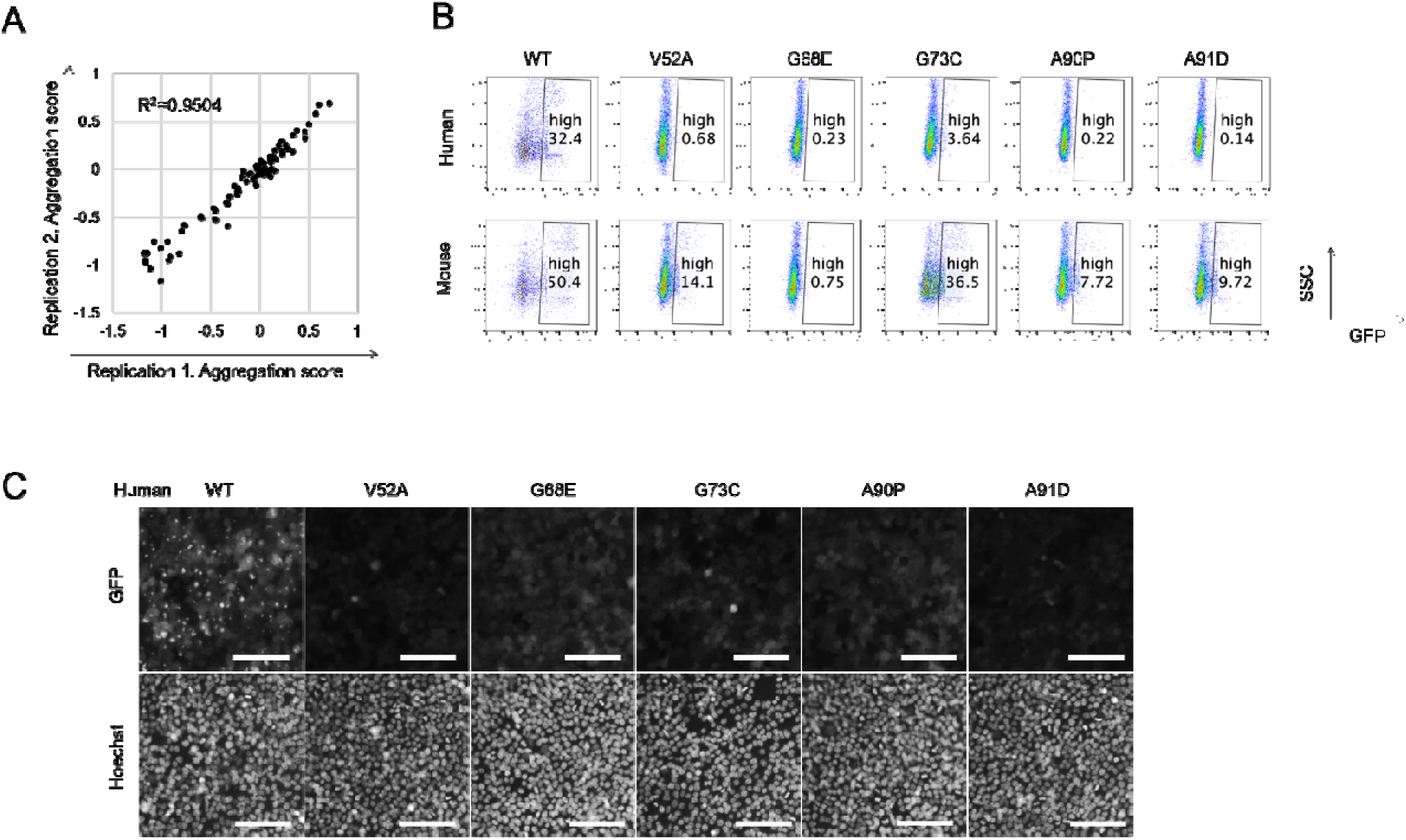
Human SNCA SNVs library screening for aggregation propensity. **(A)** The aggregation score exhibited a strong correlation between the two replicates. **(B)** The representative gating of flowcytometry to evaluate the aggregation in each mutation in the background of human and mouse α□synuclein. **(C)** Representative fluorescence microscopy images illustrating the range of α synuclein aggregation phenotypes of human variants in the background of human α□synuclein. Scale bar, 100 μm.

**Fig. S2.**
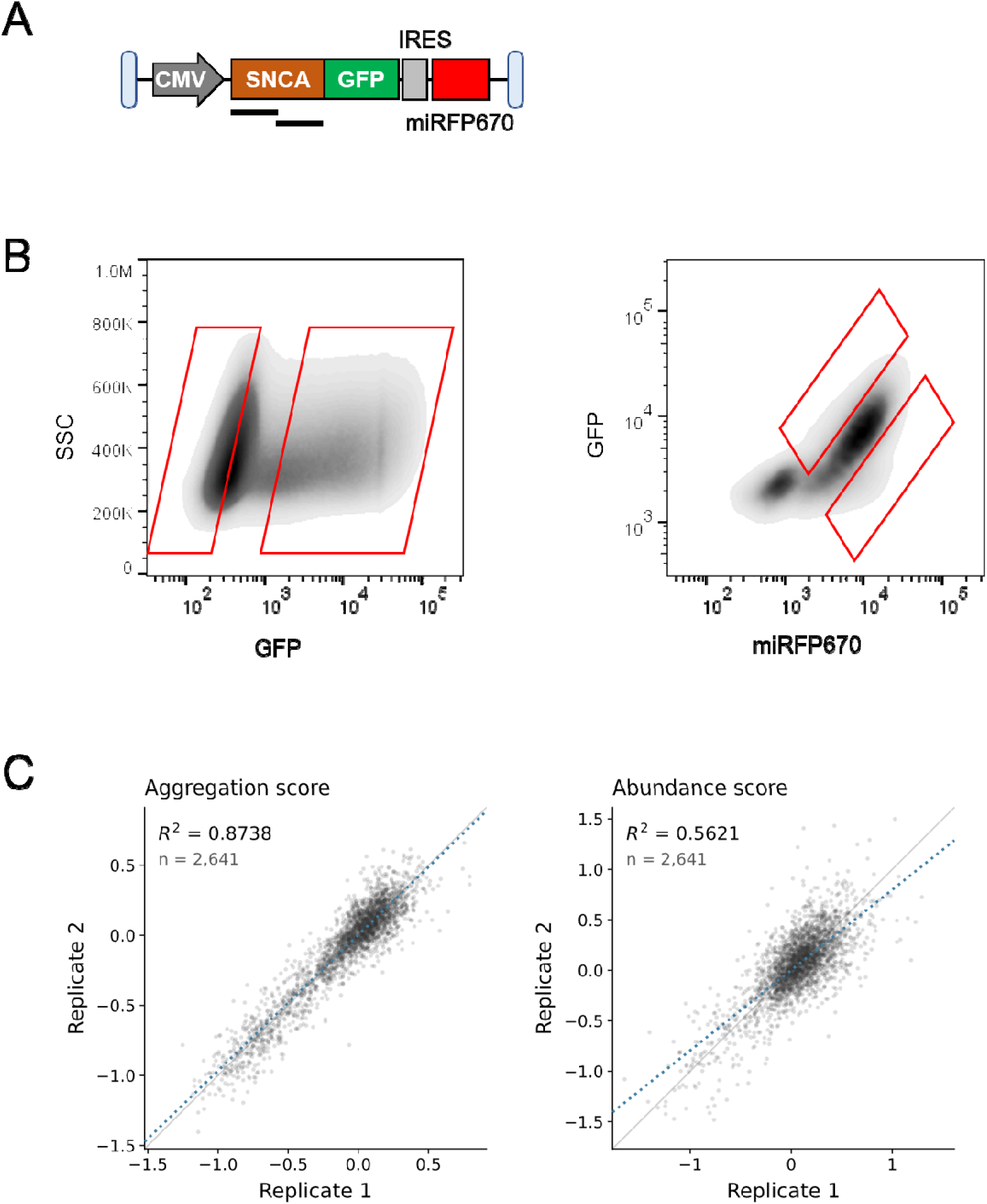
Deep mutational scanning of α□synuclein aggregation and expression. **(A)** Design of SNCA DMS library. The α□synuclein-GFP fusion protein was constructed with an internal ribosomal entry site (IRES) and miRFP670, which functions as a marker for the expression of the construct. **(B)** The gating of DMS sorting for aggregation (left) and abundance (right). **(C)** A favorable correlation was observed between the two replicates in the DMS for both aggregation (left) and abundance (right).

**Fig. S3.**
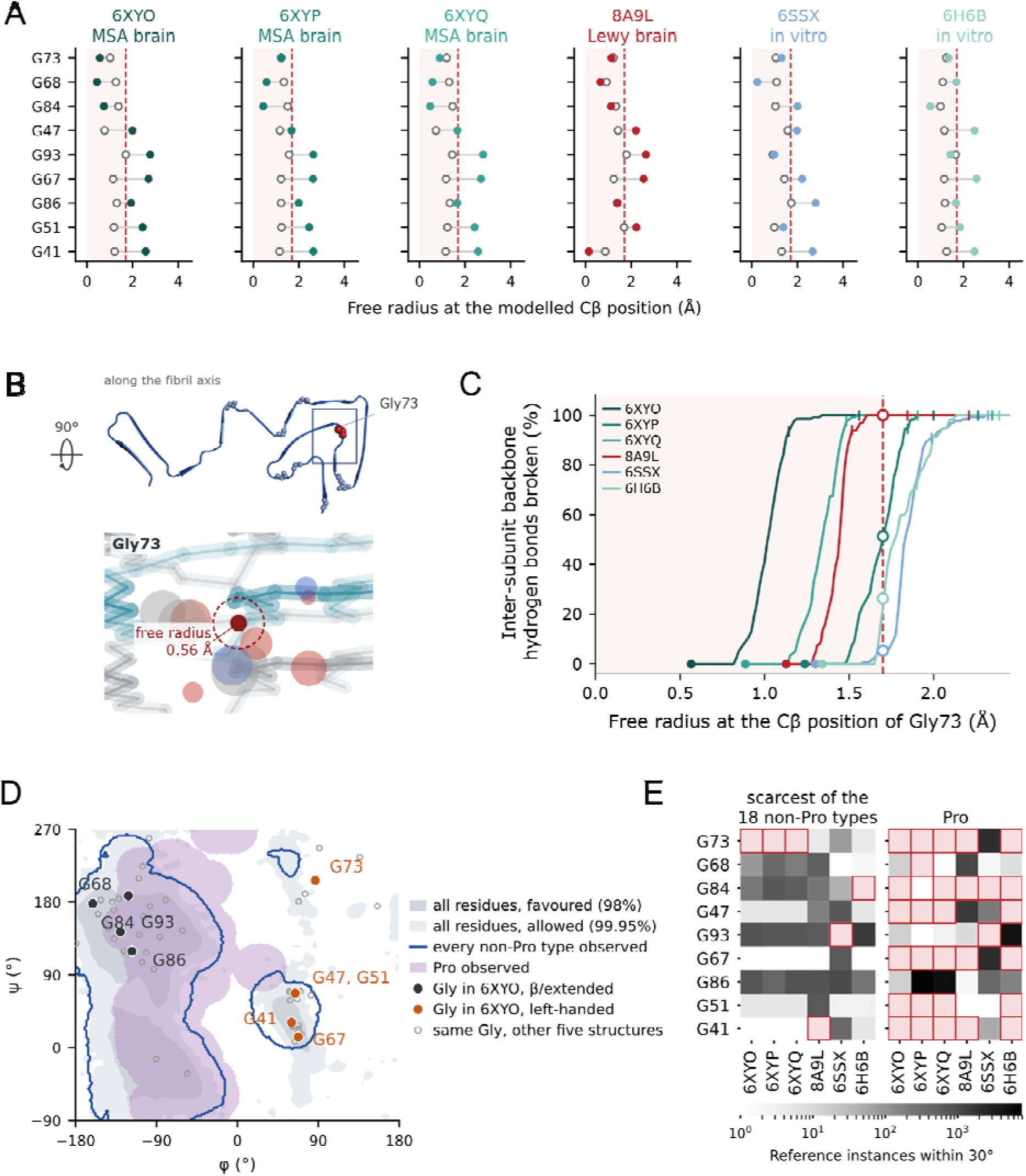
The steric criterion across six fibril structures at Gly68 and Gly73, and the backbone conformation as a negative control. (A) Free radius at the modeled Cβ position of the nine core glycines, against atoms of the same subunit (open) and of the adjacent subunits along the fibril axis (filled), in three multiple system atrophy filaments (6XYO, 6XYP, 6XYQ), one filament with the Lewy fold (8A9L) and two polymorphs assembled in vitro (6SSX, 6H6B); dashed line, carbon radius (1.70 Å). (B) Top, the 6XYO subunit viewed along the fibril axis, with Gly73 red and the other core glycines pale blue; the box marks the region drawn below and the glyph the 90° rotation between the two views. Bottom, lattice cross-section at the modeled Cβ position of Gly73, drawn as in Fig. 4D; dark red disc, the sphere that fits (0.56 Å); dashed circle, carbon. **(C)** Cost of making room at Gly73, as in Fig. 4E: the percentage of inter-subunit backbone hydrogen bonds exceeding 3.5 Å (n = 57 to 137 per structure) against the free radius reached under rigid axial expansion. Filled circles, observed rise; ticks, rises of 5.5, 6.0 and 6.5 Å; open circles, the rise at which a carbon fits (5.15 to 6.19 Å). The free radius at Gly73 is below the carbon radius in all six structures (0.56 to 1.34 Å), but expansion to the point where a carbon fits breaks every inter-subunit backbone hydrogen bond only in 6XYO, 6XYQ and 8A9L, against 51% in 6XYP, 26% in 6H6B and 5% in 6SSX. **(D)** Backbone conformation of the same positions in all six structures, against the preferences of the residues that could replace them, from 287,671 reference residues in 1,400 X-ray structures at 1.6 Å or better (glycine, proline and pre-proline excluded). Shading, 98% and 99.95% of that set; blue outline, the region in which all 18 non-proline types are observed within 30°; purple, the same for proline. Filled symbols, the glycines of 6XYO (black, β/extended; orange, left-handed); open symbols, the other five structures. **(E)** Reference instances within 30° of each of the 54 conformations, for the scarcest of the 18 non-proline types (left) and for proline (right); columns the six structures, cell shade the count on the logarithmic scale below, red cell a type not observed at all. A non-proline type is missing only at Gly73 in the multiple system atrophy filaments, Gly41 in 8A9L, Gly84 in 6H6B and Gly93 in 6SSX, while at Gly68 every type is present in every structure (1 to 532 instances); proline is unobserved in at least four structures at six of the nine positions. With 48 of the 54 conformations inside the all-18 region, conformation excludes nothing but proline at most positions and is not constant for one position across folds.

**Fig. S4.**
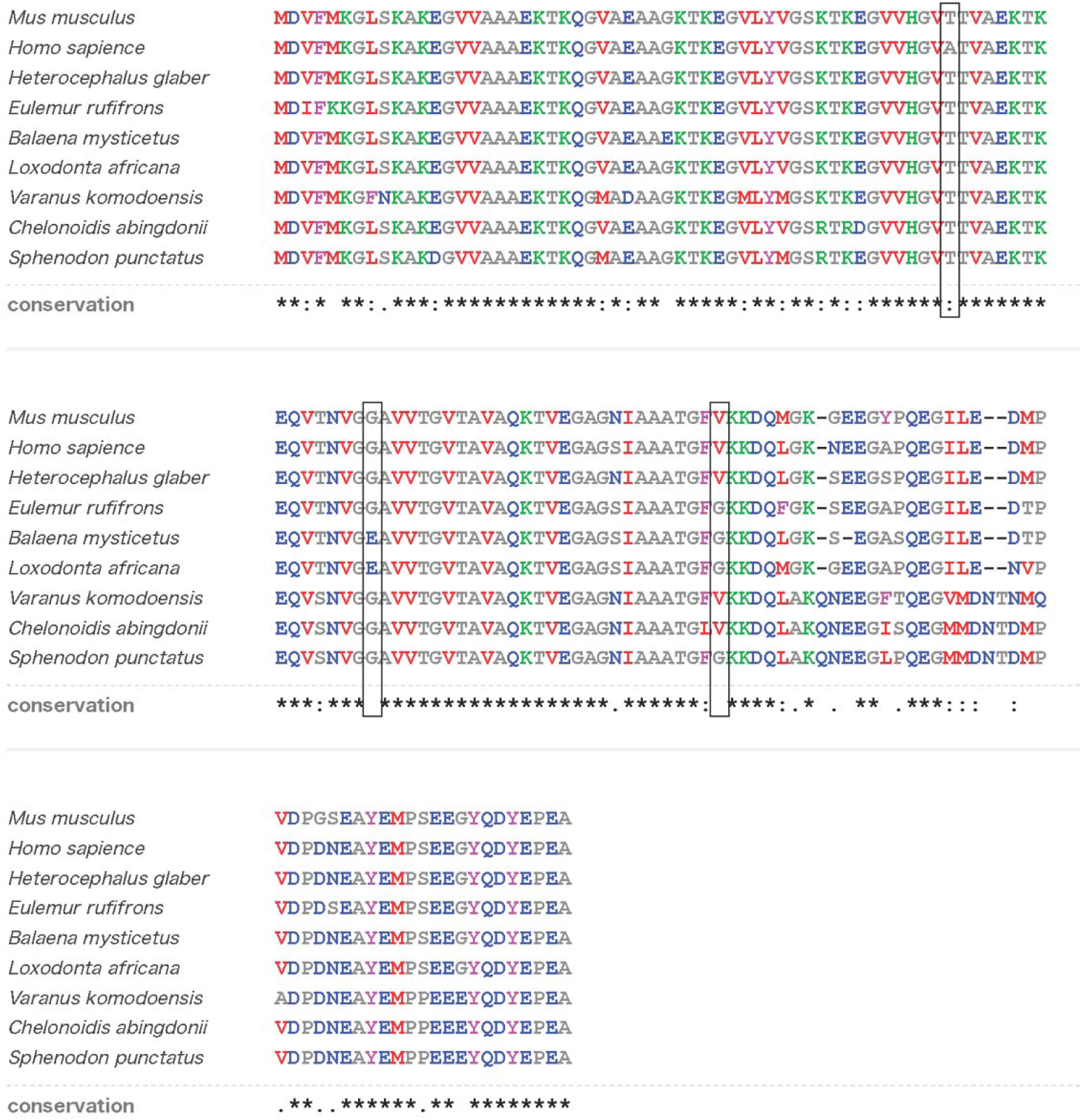
Multiple sequence alignment of α synuclein orthologs from representative vertebrate species. Amino□acid residues are colored by biochemical class, and consensus symbols (*,:,.) indicate fully conserved, strongly conserved, and weakly conserved positions, respectively. Species included span mammals, reptiles, and other vertebrate lineages, illustrating both deeply conserved motifs and lineage□specific divergence.

**Fig. S5.**
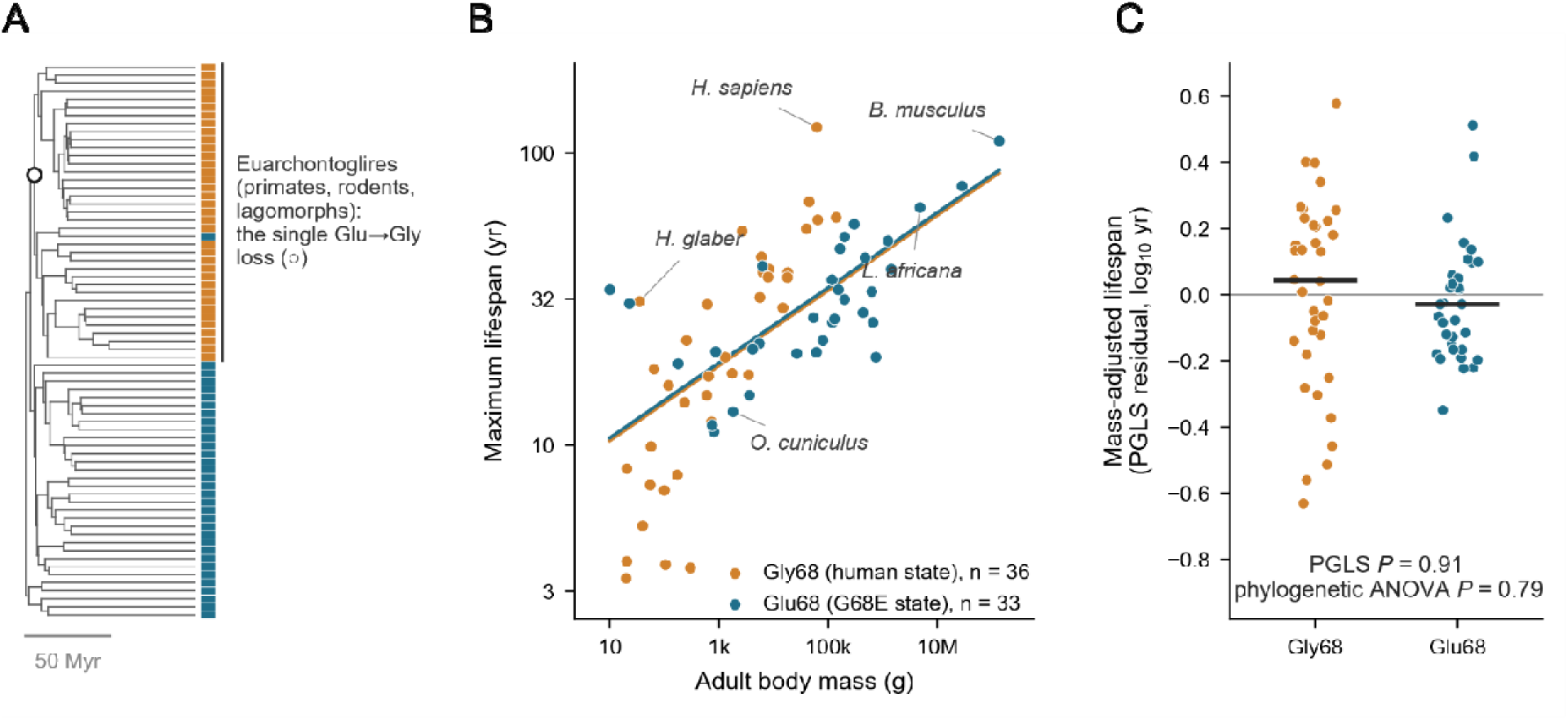
Glu68 is the ancestral placental state and is not associated with maximum lifespan after correction for phylogeny and body mass. (A) Maximum-parsimony reconstruction of residue 68 across 69 placental mammals. Tip colors give the observed state (orange, Gly68, the human state; blue, Glu68, the G68E state); the tree is the Ensembl Compara vertebrate species tree pruned to these species, made ultrametric and scaled to a 105-Myr placental root. The reconstruction requires only two changes: a single Glu-to-Gly loss on the stem of Euarchontoglires (open circle) and a single Gly-to-Glu reversal in Oryctolagus cuniculus. The same events are recovered from the 89-sequence α-synuclein alignment of Fig. S4(52 of its taxa map to tips of the species tree), so the result does not depend on species sampling. **(B)** Maximum lifespan against adult body mass for the same species, colored as in a. Lines are the phylogenetic generalized least-squares (PGLS) fit with a common slope; labeled species are Balaenoptera musculus, Homo sapiens, Heterocephalus glaber, Loxodonta africana and Oryctolagus cuniculus. **(C)** Residuals of the PGLS regression of log10 maximum lifespan on log10 adult body mass, by residue-68 state; bars are medians. Glu68 and Gly68 species do not differ (PGLS β = 0.010 log10 yr, s.e.m. = 0.084, P = 0.91; phylogenetic ANOVA P = 0.79), whereas body mass itself is strongly predictive (β = 0.129 log10 yr per log10 g, s.e.m. = 0.020, P = 2.8 × 10-8). Maximum longevity and adult body mass are from AnAge build 15.

**Fig. S6.**
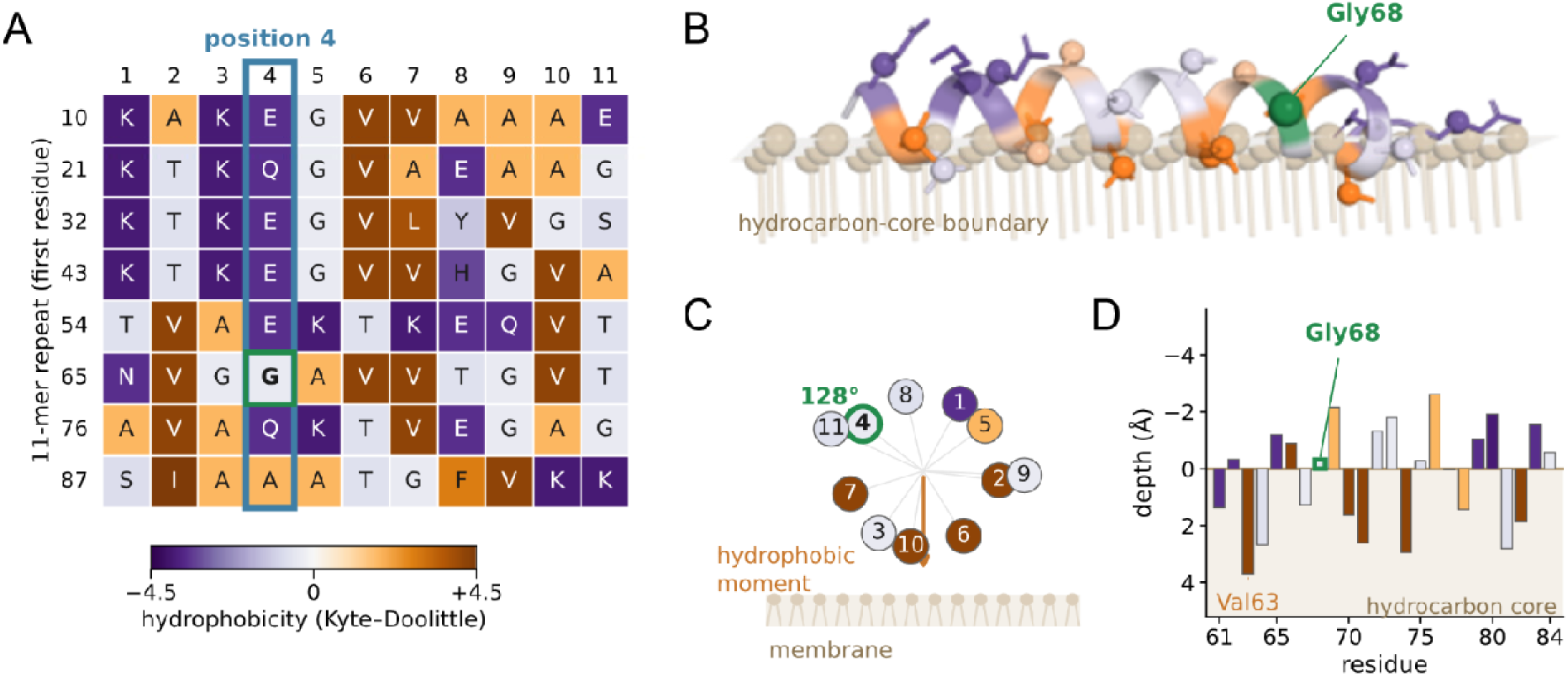
Gly68 occupies the solvent-facing position of the 11-mer repeat at which the other repeats carry glutamate or glutamine. **(A)** The eight 11-mer KTKEGV repeats (rows, labeled by their first residue) aligned by register position and colored by Kyte–Doolittle hydrophobicity. Position 4 (blue outline) carries glutamate in four repeats and glutamine in two; Ala90 and Gly68 (green outline) are the exceptions. **(B)** Residues 61 to 84 of the micelle-bound structure (PDB 1XQ8, model 1) positioned in an implicit DOPC bilayer with PPM 3.0, viewed along the membrane plane. The plane is the calculated hydrocarbon-core boundary; the lipids are schematic and drawn only behind the helix. Side chains are colored as in a; Gly68, which has no Cβ, is marked in green at its Cα. **(C)** Side-chain directions of residues 65 to 75 projected down the helix axis and labeled by register position, with the membrane below and the arrow the hydrophobic moment of residues 61 to 84. Position 4 lies 128° from the membrane direction. Glycine Cβ positions were built from the backbone; nodes closer than 22° are drawn at two radii. **(D)** Depth of the deepest atom of each residue relative to the boundary in b, colored as in a, membrane below. Eight residues insert a side-chain carbon 1.4 to 3.7 Å into the core, Val63 deepest; Gly67 and Glu61 reach it only with backbone atoms. Gly68 (green) lies 0.3 Å outside.

**Fig. S7.**
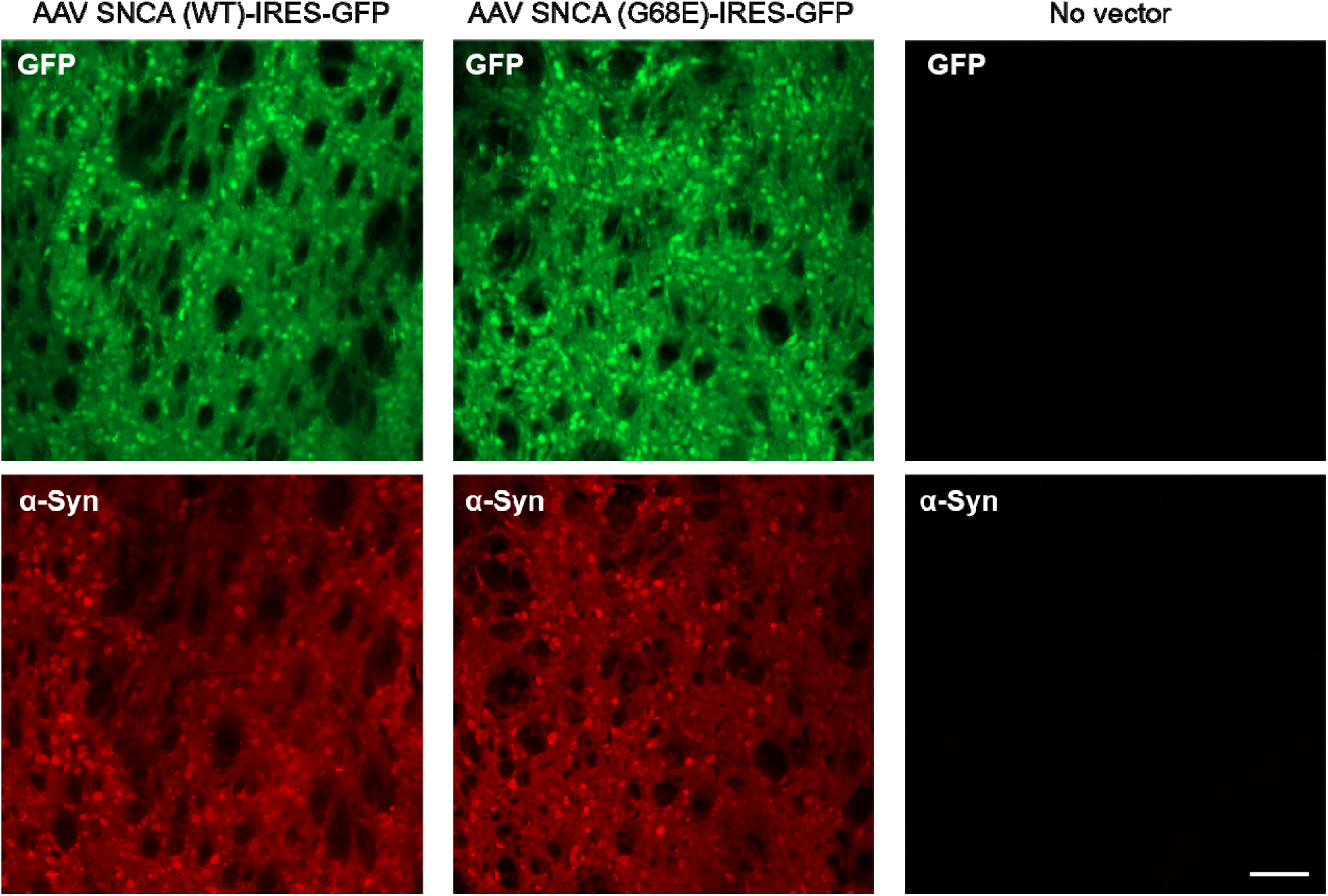
The α□synuclein expression with AAV injection in the brain of SNCA knockout mice. Three months after AAV injection, α□synuclein expression was confirmed to coincide with GFP expression. Scale bar, 100 μm.

**Fig. S8.**
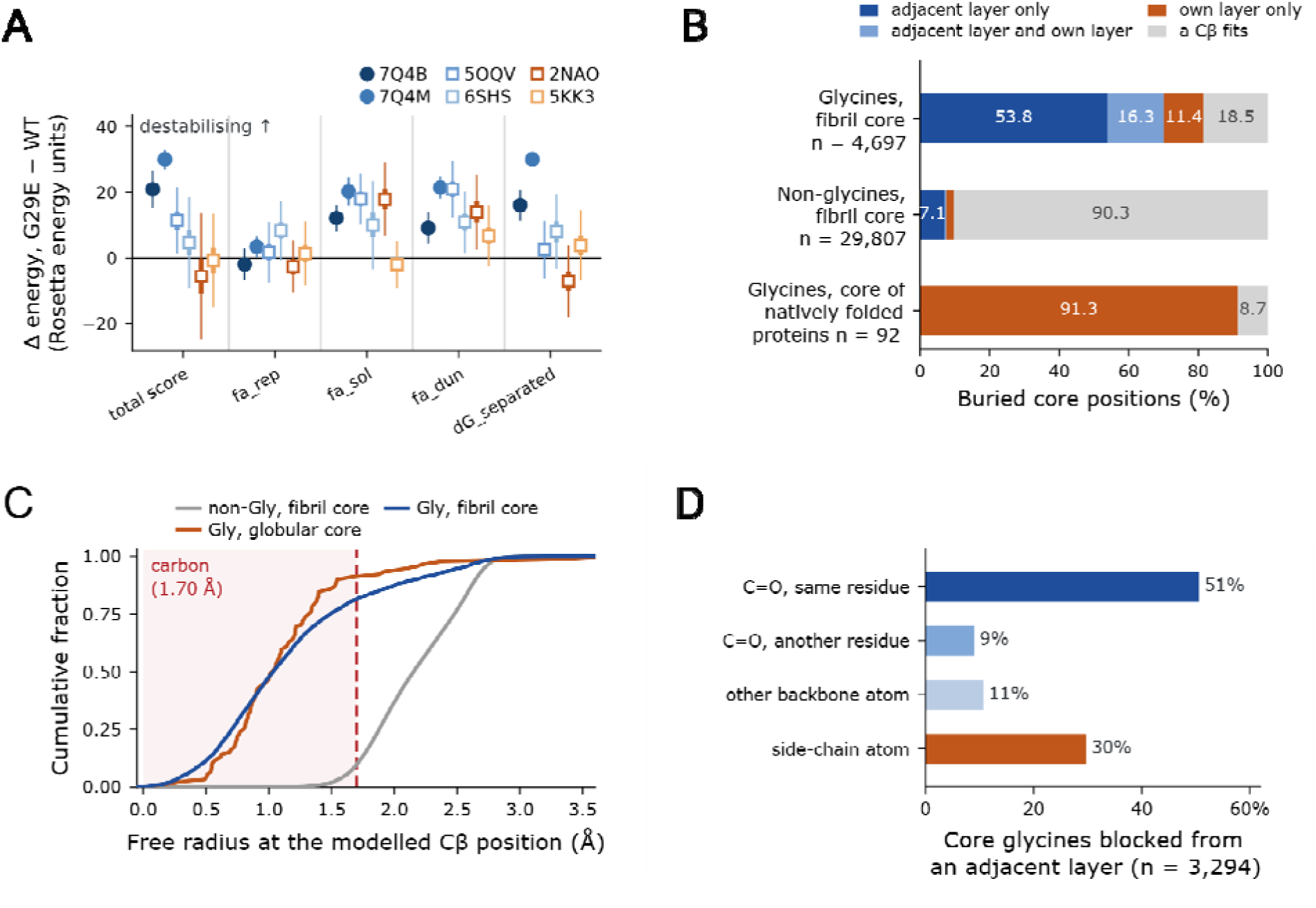
Supporting analyses for the cross-amyloid survey and for amyloid-β. **(A)** Origin of the blocking atom at buried positions (relative solvent accessibility below 0.25, one observation per structural position) for the three populations compared in the text: fibril core glycines (n = 4,697), fibril core non-glycines (n = 29,807) and buried glycines of the natively folded control set (21 structures, 18 of which contain one; n = 92). **(B)** Cumulative distributions of the free radius for the three populations. Shading and the dashed line mark 1.70 Å, the van der Waals radius of carbon. **(C)** Identity of the blocking atom for the 3,294 core glycines blocked from an adjacent layer: the backbone accounts for 70.3% and the carbonyl oxygen of the same residue one layer away for 50.6%, so the excluding atom is usually part of the hydrogen-bonded ladder that defines the cross-β architecture; the remaining 29.7% are side chains, where repacking can relieve the clash. **(D)** Rosetta term decomposition of the G29E calculation in Fig. 7D. The difference in each energy term between G29E and WT, one marker per fold, with the 95% confidence interval of the mean difference (thick) inside the standard deviation of the difference (thin), over 50 relaxed models per state. Filled circles, folds from Alzheimer brain; open squares, folds assembled in vitro or determined by NMR. In the three Aβ42 folds blocked by an adjacent layer (7Q4B, 7Q4M, 5OQV) the penalty is carried by fa_sol (12.1 to 20.2 REU) and fa_dun (9.1 to 21.4) while fa_rep stays between −2.0 and +3.4, that is by desolvation and rotamer strain rather than by raw atomic overlap; in the Aβ40 fold 6SHS, whose adjacent-layer free radius is the largest of the blocked folds (1.39 Å), fa_rep contributes comparably (+8.2) to fa_sol (+9.9) and fa_dun (+10.9).

